# Discovery of a molecular glue that enhances UPR^mt^ to restore proteostasis *via* TRKA-GRB2-EVI1-CRLS1 axis

**DOI:** 10.1101/2021.02.17.431525

**Authors:** Li-Feng-Rong Qi, Cheng Qian, Shuai Liu, Chao Peng, Mu Zhang, Peng Yang, Ping Wu, Ping Li, Xiaojun Xu

## Abstract

Lowering proteotoxicity is a potentially powerful approach for the treatment of neurological disorders, such as Parkinson’s disease. The unfolded protein response (UPR) is a major mechanism that preserves the network maintaining cellular proteostasis. In the present study, we developed the screening strategy to discover compounds that significantly enhanced the activation of mitochondrial UPR (UPR^mt^) through increasing cardiolipin content. We identified that ginsenoside Rg3 (Rg3) increased cardiolipin depending on cardiolipin synthase 1 (CRLS1) in both worms and in human neural cells. Using LiP-SMap (limited proteolysis-mass spectrometry) strategy, we identified GRB2 (growth factor receptor bound protein 2) as a direct target of Rg3 in human neural cells. Rg3 enhances the binding between GRB2 and TRKA, that transduces signals *via* phosphrorylation of ERK. We provide bioinformatic and experimental evidence that EVI1, the critical oncogenic transcriptional regulator in leukemia, binds to *CRLS1* promoter region and stimulated *CRLS1* expression and subsequently increased cardiolipin content in the presence of Rg3. In a Parkinson’s disease mouse model, Rg3 restores motor function by protecting nigral dopaminergic neurons dependent on Grb2. Our data recapitulate the TRKA-GRB2-EVI1-CRLS1 axis in maintaining proteostasis in Parkinson’s disease *via* UPR^mt^.

## Introduction

Parkinson’s disease (PD) is a neurodegenerative disease characterized by a progressive loss of dopaminergic neurons from the nigrostriatal pathway, formation of Lewy bodies, and microgliosis [1]. During the pathogenesis of PD, α-synuclein will misfold and aggregate in large quantities, forming amyloid oligomers, ribbons and fibers, which have been shown to cause neuronal toxicity by impairing numerous cellular processes [2]. Wealth of experimental data supports the hypothesis that the neurotoxicity of α-synuclein oligomers is intrinsically linked with their ability to interact with, and disrupt, biological membranes, especially those membranes having negatively-charged surfaces and/or lipid packing defects [3, 4].

Genetic data from familial and sporadic cases was used in an unbiased approach to build a molecular landscape for PD, revealing lipids as central players in this disease [5]. It is worth noting that compounds that cause fat accumulation can promote unfolded protein response (UPR) and help restore misfolded proteins [6]. Cardiolipin is a mitochondrial phospholipid involved in mitochondrial dynamics and functions [7]. Increasing the content of cardiolipin leads to stimulation of the mitochondrial UPR (UPR^mt^), which helps to maintain proteostasis, protecting neuronal cells from the accumulation of unfoled and aggregates of toxic proteins, such as amyloid-β [8]. We sought to pharmacologically manipulate UPR^mt^ that restores cellular proteostasis to treat PD.

### Rg3 improves Parkinson’s symptoms based on lipid-increasing effect and unfolded protein response

It is worth noting that fat accumulation can promotes UPR and help cells remove misfolded proteins [6]. We therefore design a quaternary screening strategy to identify lipid modulating compounds that restores cellular proteostasis to treat PD. First, we used Nile Red to stain the lipid droplet of wild type *C.elegans* N2 strain [9]. From an in-house compound library composed of over 400 natural products [10], 18 lipid-increasing compounds were found (fig. S1A-B) in the primary screening. In the BZ555 strain (*dat-1*p::GFP), GFP is expressed specifically in dopaminergic (DA) neurons under the control of dopamine transporter gene (*dat-1*) core promoter [9]. Upon 6-Hydroxydopamine (6-OHDA) treatment, DA neurons (GFP positive cells) die gradually, leading to the decrease of both GFP signal and GFP positive cell number. Using this assay, 6 compounds were found to protect DA neurons (fig. S1C-D) in the secondary screening. It has been reported that cardiolipin is necessary for the mitochondrial-to-cytosolic stress response (MCSR) induction and protects misfolded alpha-synuclein induced neural death [11]. We then check whether there are compounds that specifically increase cardiolipin content using 10-nonyl Acridine Orange (NAO) fluorescence dye [12]. We observed 3 compounds that increased NAO staining signals in the N2 strain (fig. S1E-F) in the tertiary screening. *Hsp-6* encodes a nuclear-encoded mitochondrion-specific chaperone related with the DnaK/HSP70 superfamily of molecular chaperones [6]. Using transgenic *C.elegans* SJ4100 strain (*hsp-6*p::GFP) [13], we could observe mitochondrial UPR (UPR^mt^) directly by measuring GFP signals. In the quaternary screen, we found that only ginsenoside Rg3 upregulated *hsp-6* signals (fig. S1G-I).

Further, the screening results were verified using the transgenic *C.elegans* BZ555 strain. Rg3 decreased the loss of dopamine neurons in BZ555 strain dose dependengly (Fig. 1A, fig. S2A). To evaluated the efficacy of Rg3 in removing α-synuclein aggregates, we used the transgenic *C.elegans* NL5901 strain (*unc-54p*::α-synuclein::YFP) that overexpressed α-synuclein fused with YFP (yellow fluorescence protein) [9]. The content of α-synuclein was measured by YFP signals. It turned out that Rg3 treatment reduced the accumulation of α-synuclein in NL5901 strain in a dose dependent manner (Fig. 1B, fig. S2B). Rg3 treatment also increased cardiolipin content in N2 strain (Fig. 1C, fig. S2C). Meanwhile, Rg3 dose-dependently extended lifespan (mean = 23.9 days) to that of the solvent controls (mean = 17.3 days), comparable to L-DOPA (fig. S2D-E). These data suggest that Rg3 increases cardiolipin content to stimulate UPR^mt^ and is a potent anti-AD compound.

**Figure 1.**
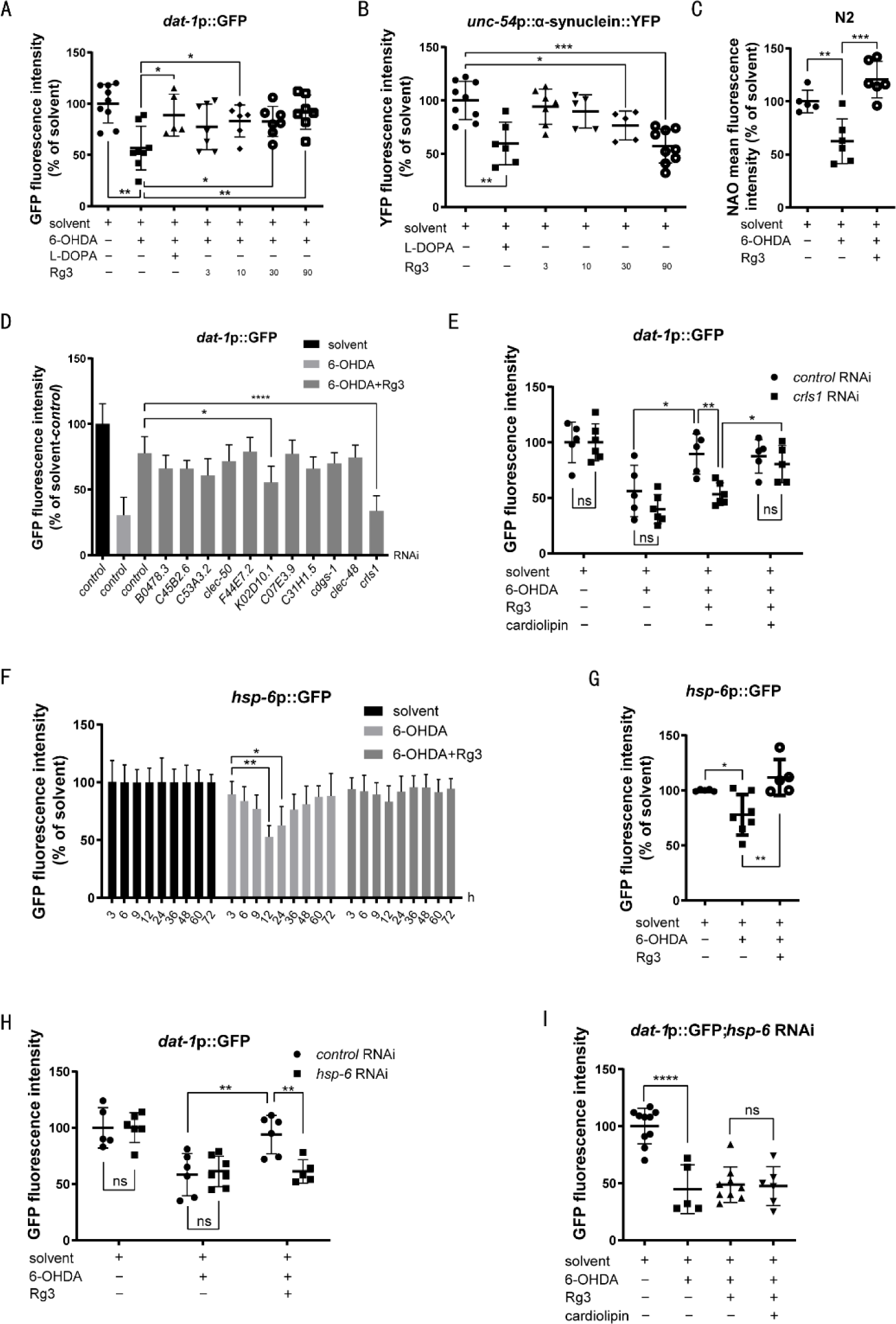
Rg3 improves Parkinson’s symptoms based on lipid-increasing effect and unfolded protein response. **(A)** The BZ555 strain was pretreated with 6-OHDA (30 mM) for 1 h then transferred to the plates with 5-Fluorouridine (0.04 mg/mL) and indicated concentrations of Rg3 or L-DOPA (10 µM) for 72 h. The fluorescence intensity of dopaminergic neurons was measured. **(B)** The eggs of NL5901 strain were treated with indicated concentrations of Rg3 or L-DOPA (10 µM). After growing to the L4 stage, the nematodes were treated with 5-Fluorouridine (0.04 mg/mL) for 12 days. The fluorescence intensity of α-synuclein protein in nematodes was measured. **(C)** The N2 strain was pretreated 6-OHDA (30 mM) for 1 h then transferred to the plates with Rg3 (90 µM) for 36h. The cardiolipin content was measured by NAO (10 μM) fluorescence signaling. **(D-E)** The BZ555 strain was cultured for three generations on culture plates with bacteria containing the indicated RNAi in the presence of Rg3 (90 µM). **(D)** Then the fluorescence intensity of dopaminergic neurons was measured. **(E)** The BZ555 strain was pretreated with 6-OHDA (30 mM) for 1 h then transferred to the plates with Rg3 (90 µM), 5-Fluorouridine (0.04 mg/mL) and cardiolipin (100 µg/ml) as indicated for 72 h. **(F)** The SJ4100 strain was treated with 6-OHDA (30 mM), 5-Fluorouridine (0.04 mg/mL) and Rg3 (90 µM) as indicated for different durations. The fluorescence representing UPR^mt^ was measured. **(G)** The SJ4100 strain was treated with Rg3 (90 µM) and 6-OHDA (30 mM) as indicated for 12 h, the fluorescence representing UPR^mt^ was measured. **(H)** The BZ555 strain fed with *hsp-6* RNAi was pretreated with 6-OHDA (30 mM) for 1 h then transferred to the plates with 5-Fluorouridine (0.04 mg/mL) and Rg3 (90 µM) as indicated for 72 h. **(I)** The BZ555 strain fed with *hsp-6* RNAi was pretreated with 6-OHDA (30 mM) for 1 h then transferred to the plates with 5-Fluorouridine (0.04 mg/mL), Rg3 (90 µM) and cardiolipin (100 µg/ml) as indicated for 72 h. * *p* < 0.05, ** *p* < 0.01, \*\*\**p* < 0.001 vs. indicated control (n=5-12).

Cardiolipin is generated and metabolized by a series of enzymes (fig. S3A), to figure out which enzyme plays a key role in maintaining cardiolipin content in PD model, each of these enzymes was knocked down by RNAi. Only *crls1* RNAi led to completely elimination of Rg3 efficacy in BZ555 worms (Fig. 1D). Consistent with previous data, suppressing *crls1* decreased cardiolipin content (fig. S3B-C). The *crls1* RNAi effect in PD model could be reversed by cardiolipin replenishment (Fig. 1E). 6-OHDA caused decrease of cardiolipin content (fig. S1E) and UPR^mt^ decrease (Fig. 1F), but did not affect ER UPR (UPR^ER^) or heat shock response (HSR), as 6-OHDA did not affect GFP signals in worm strains representing UPR^ER^ (SJ4500, *hsp-4*p::GFP) and HSR (CL2070, *hsp-16.2*p::GFP) [6], respectively (fig. S3D-E). Rg3 increased *crls1* expression (fig. S3F) might compensate the cardiolipin loss upon 6-OHDA treatment and restore UPR^mt^ (Fig. 1G). The anti-PD effect of Rg3 was abrogated by *hsp-6* RNAi (Fig. 1H). These results indicate that the anti-PD effect of Rg3 is dependent on UPR^mt^. In addition, cardiolipin supplement failed to rescue 6-OHDA treated BZ555 worms in presence of Rg3 after diminishing UPR^mt^ using *hsp-6* RNAi (Fig. 1I). Taken together, these results indicate that crls1 works upstream of UPR^mt^ in protecting 6-OHDA-induced neuron loss.

### Rg3 promotes GRB2 and TRKA binding and facilitates CRLS1 expression in SH-SY5Y cells

Next, we validated the effect of Rg3 in mammalian neural cells. Rg3 dose dependently increased cell viability after SH-SY5Y cells were exposed to 6-OHDA (Fig. 2A). HEK293 cells overexpressing GFP-tagged α-synuclein were treated with Rg3, the accumulation of α-synuclein could be easily observed by GFP fluorescence signals. Rg3 clearly reduced α-synuclein-GFP signals, comparable to L-DOPA (fig. S4A). Consistent to *C.elegans* results, Rg3 restored 6-OHDA reduced cardiolipin content in neural cells, as shown by NAO staining (fig. S4B). In mammalian cells, the induction of UPR^mt^ resulted in upregulation of mitochondrial chaperone HSP70 (heat shock protein 70) [14]. The declined HSP70 level by 6-OHDA treatment was recovered by Rg3 dose dependently (fig. S4C), indicative of increased UPR^mt^. The above data highlight the cross-species conservation of Rg3 efficacy in protecting neurons from 6-OHDA damages and α-synuclein proteotoxicity.

**Figure 2.**
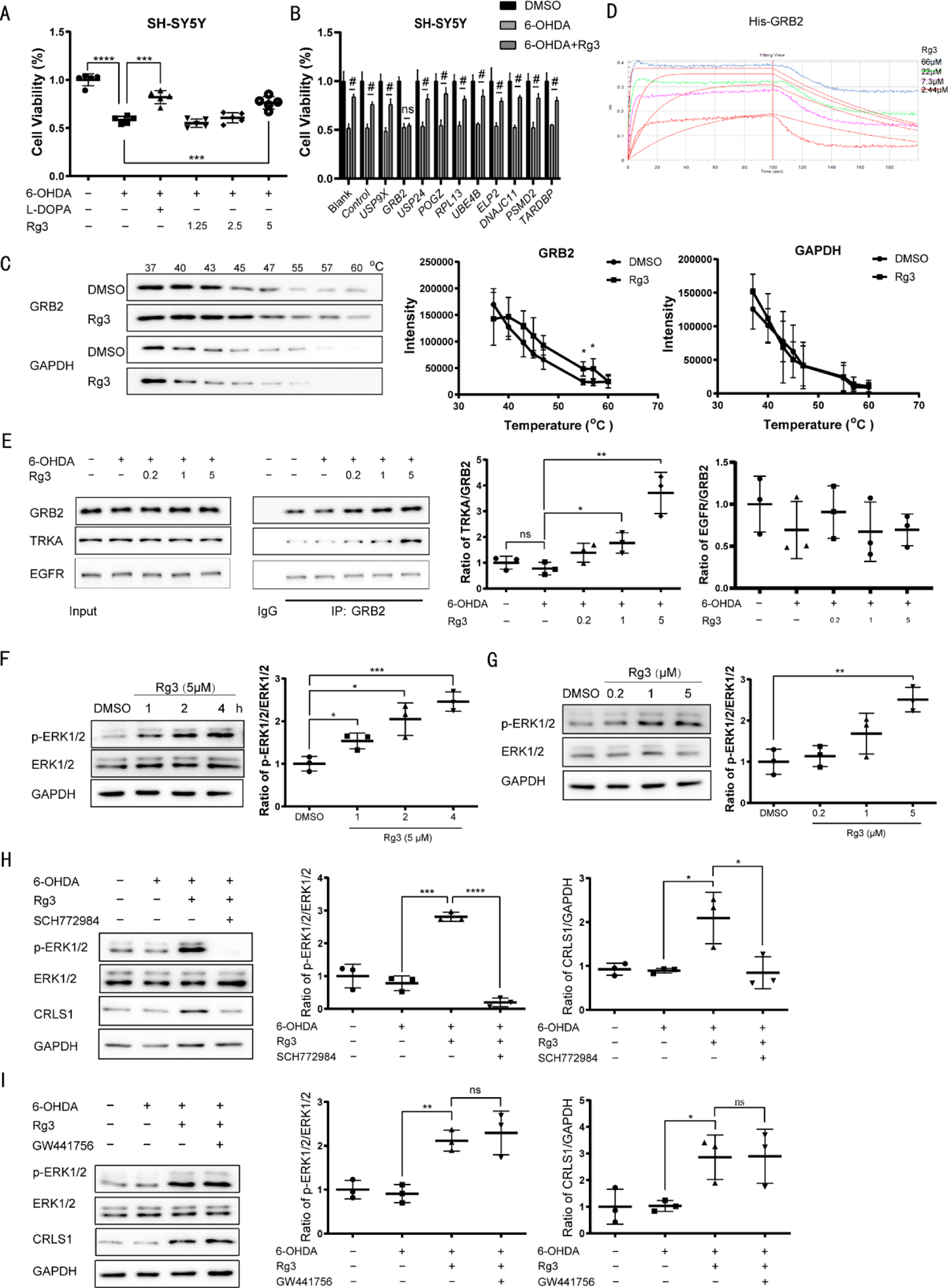
Rg3 promotes GRB2 and TRKA binding and facilitates cardiolipin synthase expression in SH-SY5Y cells. **(A)** SH-SY5Y cells were treated 6-OHDA (60μM) and indicated concentrations of Rg3 and L-DOPA (10 μM) for 24 h, then cell viability was measured by CCK8 assay. **(B)** SH-SY5Y cells were transfected with indicated siRNA for 6 h, then cultured with normal medium for 24h and treated with 6-OHDA (60μM) and Rg3 (5 μM) for additional 24 h. Cell viability was then measured by CCK8 assay. **(C)** SH-SY5Y cells were pretreated by DMSO or 5 μM Rg3 for 6 h. Then protein samples of each group were collected and applied for thermal shift assay. **(D)** Four appropriate gradient concentrations of Rg3 (2.44, 7.3, 22, 66 µM) were selected, and the proper concentrations of the GRB2 proteins were all set at 1 mg/ml for the loading solution. The kinetic parameters, including *K_on_*, *K_off_* and *K_D_* were calculated in a Fortebio Octet RED9696 instrument. **(E)** SH-SY5Y cells were treated with Rg3 (5 μM) and 6-OHDA (60μM) for 3 h. Total protein was extracted and subjected to IP study with indicated antibodies. **(F-I)** SH-SY5Y cells were treated with 6-OHDA (60μM) and Rg3 with various inhibitors, total protein lysates were collected and specific protein levels were detected by western blot. **(F)** SH-SY5Y cells were treated with 6-OHDA (60μM) and Rg3 (5 μM) for indicated periods of time. **(G)** SH-SY5Y cells were treated with Rg3 at indicated concentrations for 6 h. **(H)** SH-SY5Y cells were treated with 6-OHDA (60μM) and 0.1% DMSO, Rg3 (5 μM) or Rg3 (5 μM) + SCH772984 (10 μM) for 6 h. **(I)** SH-SY5Y cells were treated with 6-OHDA (60μM) and 0.1% DMSO, Rg3 (5 μM) or Rg3 (5 μM) + GW441756 (10 μM) for 3 h. All Experiments were repeated at least three times. \**p* < 0.05, \*\**p* < 0.01, \*\*\**p* < 0.001, #*p* < 0.001 vs. indicated control.

We then employed limited proteolysis-small molecule mapping strategy (LiP-SMap) to identify the direct target that interacts with Rg3 in proteomes [15]. We got 253 candidate proteins by LiP-SMap analysis (fig. S5A, table. S1). Ingenuity pathway analysis revealed the functional clusters of these 253 proteins (fig. S5B). According to the analysis and the literature study, we reduced the number of candidates to 10 proteins associated with Protein Ubiquitination Pathway and EIF2 Signaling, including USP9X, GRB2, USP24, POGZ, RPL13, UBE4B, ELP2, DNAJC11, PSMD2 and TARDBP. SH-SY5Y cells were transfected with siRNA silencing these candidate genes (fig. S5C, table. S2-4) and then subjected to 6-OHDA and Rg3 treatment. It turned out that only when *GRB2* was silenced, the protective effects of Rg3 against 6-OHDA induced neural damages were fully abrogated (Fig. 2B). Further, we found that Rg3 enhanced the thermal stability of GRB2 (Fig. 2C). In addition, we measured the direct binding of GRB2 protein and Rg3. The *K_D_* values of Rg3 and GRB2 were quite consistent using different methods including bio-layer interferometry (BLI) (*K_D_*=1.53 ±0.11 μM, *K_ON_* =6.16 ±0.34*10^3^ M^-1^S^-1^, *K_Off_* = 9.43 ±0.43*10^-3^ S^-1^) (Fig. 2D) and microscale thermophoresis technology (MST) (*K_D_*= 2.823 ± 4.16 μM) (fig. S5D). *In silico* molecular docking showed that Rg3 binds to Arg-67, Arg-86 and Leu-120 of GRB2 SH2 domain (fig. S5E).

It has been reported that the signaling adapter, GRB2, binds directly to the neurotrophin receptor tyrosine kinase (NTRK1 or TRKA) [16] and is essential for EGFR (epidermal growth factor receptor) signaling [17]. Then we checked whether Rg3 protects SH-SY5Y cells in a TRKA-dependent or EGFR-dependent manner. When SH-SY5Y cells were incubated with 6-OHDA (60 µM) for 3 hours, there was no change in the interaction between GRB2 and TRKA. Rg3 treatment promotes the interaction of GRB2 and TRKA in a dose-dependent manner (Fig. 2E). *In silico* molecular docking showed that binding of Rg3 to GRB2 increases the number of binding sites from 9 to 11 in the optimal binding mode of GRB2 and TRKA. (fig. S5F, table. S5-6). In contrast, Rg3 has no effect on the interaction between GRB2 and EGFR (Fig. 2E). ERK (extracellular regulated protein kinases) plays an important role in lipid synthesis [18] and is a key downstream factor related to TRKA [19, 20] and GRB2 [21]. As was shown, Rg3 increased phosphorylation of ERK in time-dependent (Fig. 2F) and dose-dependent (Fig. 2G) manners in SH-SY5Y cells. An ERK-specific inhibitor SCH772984 eliminated the effects of Rg3 on CRLS1 (Fig. 2H) and cardiolipin content (fig. S5G) [22]. However, GW441756, a specific TRKA inhibitor that binds to the extracellular domain of TRKA and blocks the binding of NGF (nerve growth factor) to TRKA (fig. S5H) [23], did not affect the Rg3 effects of activating ERK, increasing CRLS1 and cardiolipin content (Fig. 2I, fig. S5G). Rg3 promoted a higher-molecular-mass species (MW: 250-300 kDa) containing GRB2 (MW: 25 kDa) in Native-PAGE that corresponded to the estimated combined masses of TRKA (MW: 130 kDa) dimer and GRB2, which was not abolished by GW441756 (Fig. S5I). Consistently, Rg3 increased phosphorylation of TRKA, which was also not abolished by GW441756 (Fig. S5J). Together, these results indicate that Rg3 glues GRB2 and TRKA together to activate TRKA downstream signaling and then up-regulate the expression of CRLS1. This effect is independent of NGF.

### Rg3 up-regulates *CRLS1* in an EVI1 dependent manner

To understand how *CRLS1* is transcriptionally regulated, we predicted the transcription factors (TFs) and related TFs binding sites using match tools from TRANSFAC. In the *CRLS1* promoter region, we found 4 putative TFs (table. S7). Each of these TFs was knocked down by siRNAs in SH-SY5Y cells (fig. S6A). Only *EVI1 (Ecotropic Virus Integration Site 1 Protein Homolog)* siRNA abrogated the neuroprotective effects of Rg3 in the presence of 6-OHDA (fig. S6B). *EVI1* encodes a critical oncogenic transcriptional regulator for hematopoietic stem cell proliferation [24]. The function of EVI1 in leukemia and solid tumors has been well investigated [25], while the role of EVI1 in the neural system remains unknown. We first checked the subcellular localization of EVI1 in SH-SY5Y cells by immunofluorescence staining. EVI1 was mainly localized in the nuclei (Fig. 3A). 6-OHDA treatment did not change the level of EVI1 in nuclei, while Rg3 had a significant effect of increasing nuclear EVI1, which was reversed by ERK inhibitor SCH772984 (Fig. 3A). Similar results were observed in the nuclear fraction of SH-SY5Y cells (Fig. 3B). These data suggest that EVI1 is downstream of ERK signaling. When *EVI1* was knocked down, Rg3 neither promoted *CRLS1* expression (Fig. 3C-D), nor increased cardiolipin levels (Fig. 3E) in SH-SY5Y cells. Further, we validated the binding of EVI1 to *CRLS1* promoter region (Fig. 3F).

**Figure 3.**
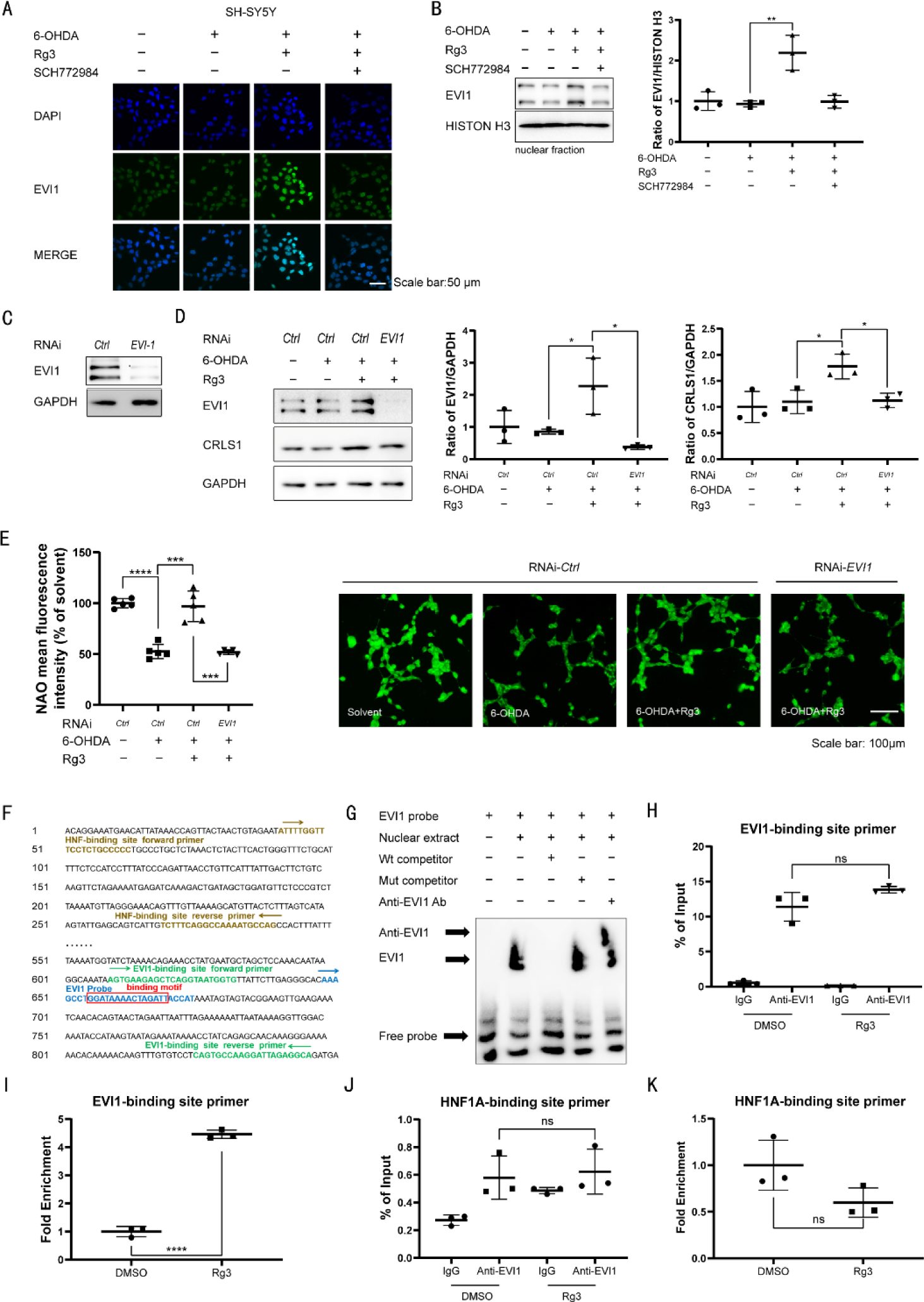
Rg3 up-regulates *CRLS1* in an EVI1 dependent manner. **(A-B)** SH-SY5Y cells were treated with 6-OHDA (60μM) and 0.1% DMSO, Rg3 (5 μM) or Rg3 (5 μM) + SCH772984 (10 μM) for 6 h. **(A)** Cells were fixed and immunostained with EVI-1 antibody. The representative images demonstrate the amount and localization of EVI-1 (green). Nuclei were counterstained with DAPI (blue). Scale bar: 50 μm. **(B)** Nucleoproteins were extracted and subjected to western blot with indicated antibodies. **(C-E)** SH-SY5Y cells were transfected with indicated siRNA for 6 h, then cultured with normal medium for 48h. Then Cells were treated with 6-OHDA (60μM) and Rg3 (5 μM) for 6 h. **(D)** Total proteins were extracted and subjected to western blot with indicated antibodies. **(E)** The cardiolipin content was measured by NAO (5 μM) fluorescence signaling. Scale bar: 100 µm. **(F)** A putative EVI1 probe binding site, a pair of EVI1 binding site primers and a pair of HNF1A binding site primers in the *CRLS1* promoter. **(G)** Electrophoretic mobility shift assay (EMSA) of EVI1 binding to the *CRLS1* promoter was performed with wildtype and mutant probes. Wt, wild type. Mut, mutant. Ab, antibody. **(H-I)** ChIP-qPCR analysis for the binding of EVI1 to the *CRLS1* promoter in SH-SY5Y cells treated with Rg3 (5 μM). **(J-K)** ChIP-qPCR analysis for the binding of HNF1A to the *CRLS1* promoter in SH-SY5Y cells treated with Rg3 (5 μM). All Experiments were repeated at least three times. \**p* < 0.05, \*\**p* < 0.01, \*\*\**p* < 0.001, \*\*\*\**p* < 0.0001 vs. indicated control.

Electrophoretic mobility shift assays (EMSA) demonstrated that the 655/668 region (GGATAAAACTAGATT) was probably the EVI1 binding motif within the fragment of the *CRLS1* promoter (Fig. 3F-G, table S8). We further performed chromatin immunoprecipitation (ChIP) assays and found that EVI1 was recruited to the 609 to 845 bp region that contains a predicted EVI1 binding motif (Fig. 3H-I, table S9). In contrast, Rg3 did not affect other transcription factor HNF1A (Fig. 3J-K, table S9). Overall, these results indicate that EVI1 is a *bona fide* transcription factor of *CRLS1*. Rg3 increases the expression of *CRLS1 via* EVI1.

### Rg3 ameliorates MPTP-induced behavioral deficits through Grb2

Motor dysfunction is the main pathological character of PD patients and mouse models [26, 27]. MPTP produced reliable and reproducible damages to the nigrostriatal dopaminergic neurons [26]. To examine the protective effects of Rg3 in PD mouse model, we first measured motor function in the open field, accelerating rotarod, and pole tests on the day after the final Rg3 or normal saline (NS) administration in MPTP-treated mice (Fig. 4A). Total travelled distance during 3 minutes in mice treated with MPTP significantly reduced compared to control mice, which indicated the defect in autonomous activity. These impairments were reduced after Rg3 administration as the distance traveled by mice increased dramatically (Fig. 4B-C). In accord with pervasive motor dysfunction, MPTP-lesioned mice required more time to descend the pole and fell off the rotarod in less time than controls. Rg3 treatment reduced pole descent time (Fig. 4D). Further, Rg3 significantly increased rotarod time compared with the solvent group (Fig. 4E). Knockdown of *Grb2* in mouse neural cells (N2a) reversed Rg3 effects against 6-OHDA damages (fig. S7A-B), consistent to the results in human neural cells (Fig. 2B). AAV5 virus [28] carrying *Grb2* shRNA silenced *Grb2* in the substantial nigra (SN) of mice (fig. S7C, table. S3-4), which reversed the behavior benefits of Rg3 (Fig. 4B-E). Consistent to cell results, Rg3 increased Crls1 expression in the substantial nigra (SN) of mice (Fig. 4F). We next examined the neuroprotective efficacy of Rg3 by comparing nigral dopaminergic neuron numbers. MPTP-lesioned mice exhibited severe loss of tyrosine hydroxylase (Th)-protein and Th-positive neurons. Rg3 treatment greatly rescued Th level and Th-positive neurons (Fig. 4F-G). These findings suggested that the motor improvements (Fig. 4B-E) resulted from protection of nigrostriatal dopaminergic neurons. MPTP-lesioned mice exhibited enhanced Il-1β levels in the SN, which was reversed by Rg3 administration (Fig. 4G). The neuroprotection and anti-inflammation effects of Rg3 was completely abolished when *Grb2* in the SN was suppressed by AAV5-sh*Grb2* transduction. Together, these results suggest that Grb2 is essential for the therapeutic effects of Rg3 in protecting nigral dopaminergic neurons in mice.

**Figure 4.**
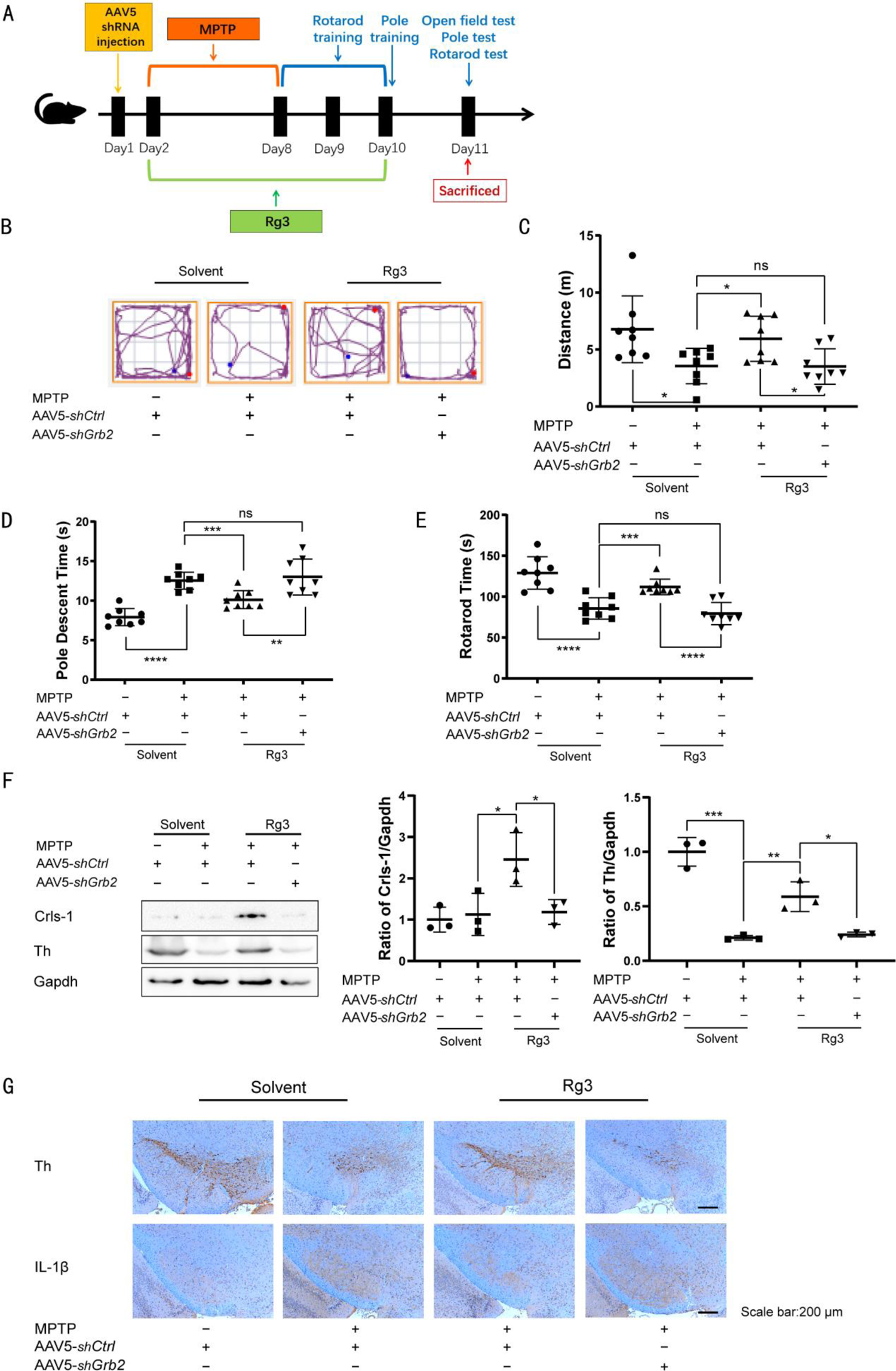
Rg3 ameliorates MPTP-induced behavioral deficits through Grb2. **(A)** The scheme of experimental procedure. A behavior-disordered PD model was induced by MPTP in C57BL/6J mice. Mice were then injected with AAV5-sh*GFP* (control group) or AAV5-sh*Grb2* (*Grb2*-interfering group) in substantia nigra (n = 8 for each group). **(B-C)** Open field analysis of indicated mice. Total travelled distance in 3 minutes during test were recorded and analyzed. **(D)** Times required for mice to descend from the pole were recorded and analyzed. **(E)** Latency to fall during test on accelerated rotarod trials was recorded and analyzed. **(F)** Immunoblotting analysis of Th and Gapdh in the SN from 4 groups of mice as indicated. **(G)** Immunohistochemical staining for Th and Il-1β in the SN in these 4 groups of mice as indicated. \**p* < 0.05, \*\**p* < 0.01, \*\*\**p* < 0.001, \*\*\*\**p* < 0.0001 vs. indicated control.

## Discussion

Brain tissue of PD patients and animal studies have shown that oxidative stress damages key cellular pathogenetic proteins that in turn cause these misfolded proteins to form small aggregates, facilitating PD development and multiple system atrophy [29, 30]. Maintaining proteostasis is critical to restore neural cell function and protect from cell death [31]. In the present study, we identified that Rg3 decreased the loss of dopamine neurons in BZ555 strain (Fig. 1A, fig. S1C-D) and promoted the cell viability in mammalian cells challenged by 6-OHDA or α-synuclein overexpression (Fig. 2A, fig. S4A). The function of Rg3 seems to be dependent on its role in enhancing cardiolipin content and UPR^mt^ that relies on a critical enzyme Crls1 controling cardiolin synthesis (Fig. 1E-I).

Ginsenoside Rg3, one of the major active components of ginseng, exhibits antitumor activity in various tumors [32] and neuroprotective effects with anti-oxidative stress and antiinflammation [33]. Although Rg3 has been shown to be neuroprotective by protecting N-Methyl-d-aspartate (NMDA)-induced neuronal death [34, 35] and inhibiting the opening of mitochondrial permeability transition pores to produce antioxidant effect in the brain [36], but the direct target proteins of Rg3 have not been identified so far. Using LiP-SMap methods, we were able to identify GRB2 as the potent Rg3 target protein in regulating cardiolipin homeostasis and relieving symptoms of PD. Of course, we select GRB2 among dozens of genes. The other genes might also interact with Rg3. However, these genes probably do not play major roles in the process of Rg3 induced neuroprotection (Fig. 2B). Mechanistically, we found that Rg3 promoted the binding of GRB2 and TRKA, thereby activating ERK phosphorylation and stimulated transcription factor EVI1 nuclear localization and then activated *CRLS1* transcription (Fig. 3A-D).

NGF is highly expressed in the human substantia nigra (SN) and decreased dramatically during PD patients and animal models [37–40]. The use of NGF to treat neurodegerative disease is limited due to its constant occurred side effects including pain response and weight loss [41]. Rg3 helps to form TRKA-GRB2 complex that transduces signaling to activate ERK even independent of NGF (Fig. 2H-I). The function of Rg3 is quite unique than other molecular glue that normally aim to degrade target proteins [42, 43]. To our knowledge, this is the first molecular glue that activates a tyrosine kinase receptor. In addition, we discovered a novel EVI1-CRLS1 pathway that support neuroprotective effects of NGF/TRKA signaling pathway, besides cAMP response element-binding protein (CREB) [44–48]. This is a novel therapeutic strategy that maybe applicable to neurodegenerative disorders caused by proteopathies (Fig. S8).

## Acknowledgements

ACKNKOWLEDGEMENTS

We thank Prof. Wenyuan Wang at the Interdisciplinary Research Center on Biology and Chemistry for help with Parkinson’s disease model and human genetics.

## Author contributions

X.X. conceived the project. L.-F.-R.Q., C.Q., S.L., and X.X. designed the experiments. L.-F.-R.Q., C.Q., S.L., C.P., M.Z., P.W., performed the experiments. L.-F.-R.Q., C.Q., S.L., C.P., M.Z., P.W., P.Y., P.L., and X.X. analyzed the data. L.-F.-R.Q., and X.X. wrote the paper with input from all authors.

## Materials and methods

### Reagents

Ginsenoside Rg3 was obtained from Chengdu Pufei De Biotech Co., Ltd (Sichuan, China), which was dissolved with 1% dimethylsulfoxide (DMSO) for the *in vitro* experiments, or with 0.5% carboxymethylcellulose sodium (CMC-Na) solution for the *in vivo* tests. 5-Fluorouridine, Acridine Orange 10-nonyl bromide (NAO), Nile Red, 1-Methyl-4-phenyl-1,2,3,6-tetrahydropyridine hydrochloride (MPTP), EZ-ChIP kit, KH_2_PO_4_, Na_2_HPO_4_, NaCl, iodoacetamide, protease K, DMSO, formic acid were purchased from Sigma-Aldrich (St. Louis, MO, USA). SCH772984 (HY-50846) and GW441756 (HY-18314) were purchased from MedChem Express (Shanghai, China). Tissue protein extraction kit, bicinchoninic acid (BCA) protein assay kit, EMSA/Gel-Shift kit, Coomassie blue G250 and DAPI (cat. No. AF0006) were obtained from Beyotime Institute of Biotechnology (Jiangsu, China). CMC-Na was purchased from Sangon Biotech Co., Ltd (Shanghai, China). AceQ qPCR SYBR Green Master Mix and HiScript QRT SuperMix for qPCR were purchased from Vazyme Biotech Co., Ltd. (Nanjing, China). Primary antibodies used in this study include Hsp70 Rabbit mAb (cat. No.AF1156, Beyotime, China), GAPDH Mouse mAb (cat. No. AF5009, Beyotime, China), TrkA Antibody (cat. No. 2505, Cell Signaling Technology, Beverly, USA), EGF Receptor Rabbit mAb (cat. No. 4267, Cell Signaling Technology, Beverly, USA), EVI-1 (C50E12) Rabbit mAb (cat. No. 2593, Cell Signaling Technology, Beverly, USA), Anti-rabbit IgG (H+L)-F(ab’)2 Fragment-Alexa Fluor 555 Conjugate (cat. No. 4413, Cell Signaling Technology, Beverly, USA), CRLS1 pAb (cat. No. PA5-25338, Thermo, USA), GRB2 Rabbit mAb (cat. No. A19059, ABclonal, China), ERK1/ERK2 Rabbit pAb (cat. No. A16686, ABclonal, China), Phospho-ERK1-T202/Y204 + ERK2-T185/Y187 Rabbit pAb (cat. No. AP0472, ABclonal, China), Phospho-TrkA-Y490 Rabbit pAb (cat. No. AP0492, ABclonal, China), TH Rabbit pAb (cat. No. A0028, ABclonal, China), Histone H3 Rabbit pAb (cat. No. A2348, ABclonal, China), IL1 beta Rabbit pAb (cat. No. A16288, ABclonal, China). Secondary antibody was purchased from Beyotime Institute of Biotechnology (Jiangsu, China). DMEM (cat. No. 11965-092) and FBS (cat. No.10100147) were from Gibco (USA). Penicillin and streptomycin, Trpsin and 4% paraformaldehyde were from KeyGEN (China). Mass spectrometry-grade trypsin was from Promega. Electrochemiluminescence (ECL) was from Tanon (China).

### Worm Strains, culture, and synchronization

Wild-type: N2, NL5901 (pkIs2386 [*unc-54*p:: α-synuclein::YFP + unc-119(+)]), BZ555 (egIs1 [*dat-1*p::GFP]), SJ4100 (zcIs13[*hsp-6*p::GFP]), SJ4500 (zcIs4 [*hsp-4*p::GFP]V), CL2070 (dvIs[*hsp-16.2*p::GFP]) were obtained from the Caenorhabditis Genetics Center (University of Minnesota). Following the standard protocol, worms were fed by the lawn of *Escherichia coli* strain OP50. The worms were grown on solid nematode growth medium (NGM) plate in 20 °C incubator. The synchronized eggs were isolated by bleaching gravid adult worms in bleaching solution (12% NaClO and 10% 1M NaOH). Then, eggs were transferred into fresh NGM plate without OP50 and incubated overnight at 20 °C to obtain synchronized L1-stage worms. [1]

### Inverted fluorescence microscope and image processing

Worms were immobilized with tetramisole and mounted on 6% agarose pads on glass slides. Images were acquired using inverted fluorescence microscope. For each condition, 5-7 fluorescence images of different worms were taken. Image processing was performed with Fiji software.

### RNAi Treatment and RNAi Screening

RNAi treatment and RNAi screening were done according to a former research [2]. Briefly, synchronized eggs were harvested by bleaching and nematodes were grown on plates with *E. coli* OP50 until they reach early adulthood before transferred to RNAi plates. Day 1 adult worms are then grown on the RNAi plates with *E. coli* HT115 that carried the RNAi construct for 3 days at 20°C unless indicated otherwise in the figure legends. Worm eggs were synchronized by bleaching and grown on *E. coli* OP50 plates until they reach day 1 adult at 25°C. Day 1 adult worms were transferred to *E. coli* HT115 RNAi plates and grown for 2 more days before the analysis.

### Lipid deposition analyses

The nematode strain N2 worms [2] were washed off from plates with M9. Briefly, worms were fixed with freshly made 0.5% paraformaldehyde and frozen in liquid nitrogen immediately. M9 with Nile Red was added (1 μg/ml in final concentration) to the worms prior to staining for 15-30 min. Using a high-content screening instrument to take the fluorescence image of each well (TTP Labtech).

### Nonyl Acridine Orange Staining

Nonyl Acridine Orange (NAO) was used to stain cardiolipin in nematode strain N2 [2]. After fixation and washing, 10 μM of NAO solution is added for 15-30 min, the fluorescence signaling represented cardiolipin content in worms.

### Analysis of dopaminergic neurodegeneration and activation of UPR

Nematode strain BZ555 was used for analysis of dopaminergic neurodegeneration [1]. Nematode strain SJ4100, SJ4500 and CL2070 were used for analysis of UPR [2]. Synchronized L3 worms were treated with 30 mM 6-OHDA [3], 10 mM ascorbic acid, and diluted OP50 solution to induce DA neuron degeneration. Synchronized L3 worms were treated with 30 mM 6-OHDA (dissolved in 10 mM ascorbic acid) and diluted OP50 solution to induce DA neuron degeneration. Induced worms were then transferred onto NGM/OP50 plates containing indicated compound and 0.04 mg/ml 5-Fluorouridine. DMSO at final concentration of 1% (v/v) was applied in all treatments and used as a vehicle control. The fluorescence intensity of dopaminergic neurons was measured using inverted fluorescence microscopy (Nikon Ts2R).

### Analysis of alpha synuclein protein aggregation

The α-synuclein expressing worms, NL5901 [1], were firstly synchronized. The newly hatched worms were transferred onto NGM/OP50 plates containing indicated compound and 0.04 mg/ml FUDR, incubated at 20 °C for 12 days.

### Lifespan tests

Lifespan analyses were performed on synchronized L4 NL5901 [1] worms treated with indicated compound. The worms were counted and classified as alive, dead, or censored every day. The data were plotted as a survival curve and compared between groups.

### Cell culture and Maintenance

SH-SY5Y cell, HEK293 and Neuro-2a (N2a) cell lines were obtained from the American Type Culture Collection. All of the cell lines were grown in DMEM (Gibco) containing 100 units/ ml penicillin and 100 μg/ml streptomycin sulfate (Keygen Biotechnology) with 10% FBS (Gibco) under 37°C, 5% CO2.

### CCK8 assay

An *in vitro* Parkinson’s disease (PD) model was established by treating SHSY5Y cells with 6-OHDA [4]. Cells were seeded in 96-well plate at a density of 1 × 10^4^ cells *per* well for 24 h. After that, cells were treated with various concentrations of compounds. The proliferation of SH-SY5Y cells was evaluated by the cell counting kit-8 (CCK8) assay according to the manufacturer’s instructions.

### Transfection

HEK293 cells were transfected with human alpha-synuclein overexpressioin plasmid (Cat: HG12093-ANG, Sinobiological, Beijing, China) using Lipo3000. Human alpha-synuclein was tagged with N-terminal GFP. After incubating with plasmids for 6 hours, the supernatant was replaced by drug-containing solution or the control solution for 42 hours incubation. The amount of alpha-synuclein in cells was imaged by fluorescence microscopy (Nikon Ts2R).

### Cell Cardiolipin Staining

SH-SY5Y cells were treated with blank solvent, 6-OHDA, and/or Rg3 for 24 hours. After 24 h of incubation, the supernatant was replaced by phenol red-free medium containing 5 μM NAO for 20 minutes on ice in the dark. NAO was washed out by phenol red-free medium 2-3 times. Fluorescence images were captured with an inverted fluorescence microscopy (Nikon Ts2R) and the average fluorescence intensity of each image was analyzed with Fiji software.

### Limited proteolysis combined with mass spectrometry (LiP-SMap)

LiP-SMap was done according to a former report [5]. Briefly, proteomes were extracted and incubated with Rg3 or DMSO for 10 minutes at 25°C. Samples were subjected to limited proteolysis to generate structure-specific protein fragments. Fragments were then digested with the sequence-specific protease trypsin to generate peptide mixtures amenable to bottom-up proteomic analysis. The polypeptide samples were dissolved in 0.1% formic acid and detected by Orbitrap mass spectrometer (Thermo), with label-free quantitation (LFQ) analysis, search for differential proteins on MaxQuant platform *via* MS sensitivity.

### Immunoprecipitation

Cells were harvested and suspended in lysis buffer (Cell Signaling Technology, 9806) containing protease and phosphatase inhibitors. The cell lysates were centrifuged for 10 min at 14,000 g at 4°C. About 10% of the supernatant was used for western blot as inputs, while the rest of homogenates were incubated with specific antibodies at 4°C overnight. Then add protein A/G sepharose beads (Thermo Fisher Scientific, 78609) for 2 h at 4°C. The immunoprecipitation beads were washed with cold PBS for five times, followed by western blotting analysis.

### RNA interference

The siRNA oligos as listed in table S3 were synthesized by BIOSYNTECH (Suzhou, China). Cells were transfected with the siRNAs using Lipofectamine™ 3000 (Invitrogen) according to the manufacturer’s manual and then harvested after 48 h. Control vector-transfected cells were used as controls for all the experiments.

### Western blot

The protein samples from the brain tissues and cultured cells were harvested and suspended in lysis buffer (Beyotime Biotechnology, P0013J), and the protein contents were determined using the BCA method (Beyotime Biotechnology, P0009). The extracted proteins were separated by SDS-PAGE and then transferred to a nitrocellulose membrane. The membranes were blocked for 1 h at room temperature and then treated overnight at 4°C with the primary antibodies. After incubated with HRP conjugated secondary antibodies for 2 h at room temperature, immunoblots of the membranes were visualized by chemiluminescence detection kit (Tanon Science & Technology). To eliminate variations due to protein quantity and quality, the data was adjusted to GAPDH (glyceraldehyde-3-phosphate dehydrogenase).

Cells were lysed with Native-PAGE lysis buffer (50 mM Tris (pH 7.4), 150 mM NaCl, 1% NP-40, 0.25% sodium deoxycholat and protease inhibitor). The protein contents were determined using the BCA method (Beyotime Biotechnology, P0009). Lysates were mixed with Native-PAGE sample buffer 5X (Beyotime) and subjected to Native-PAGE using ExpressPlus™ 4-12% PAGE Gel (GenScript) and Tris-MOPS-SDS Running Buffer (GenScript). Then extracted proteins were transferred to a nitrocellulose membrane. The blocking and visualization steps are the same as above.

### Real-time PCR

Total RNAs were isolated using Trizol reagents (Vazyme) according to the manufacturer’s instructions. RNA concentrations were equalized and reverse transcribed into cDNA using HiScript Reverse Transcriptase kit (Vazyme). Gene expression level was measured by qRT-PCR using SYBR-green (Vazyme). Gene expression was normalized to GAPDH, primers used in this study was listed in table S4.

### Immunofluorescence

Formaldehyde-fixed cells were incubated with indicated primary antibodies at 4 °C overnight, and then incubated with secondary antibodies for 1 h at room temperature. DAPI was added and incubated with cells for 5 min. Immunofluorescence was visualized and captured on a confocal microscope (Olympus FV3000).

### *In Silico* molecular docking

To analyze the binding affinities of small molecule compound to target protein and the possible binding sites, an *in silico* protein ligand docking software AutoDock 4.2 or Discovery Studio (BIOVIA) program was applied [6]. Briefly, Crystal structure file of target protein was downloaded from the RCSB protein data bank. Docked conformations were generated by Lamarckian genetic algorithm (LGA). After statistical analysis, the conformations with the lowest binding energies were performed to generate the interaction figures of ligands to target protein.

### Binding of Rg3 to GRB2

The interaction of Rg3 and recombinant GRB2 was monitored by BLI using a Fortebio Octet RED96 instrument. In the gradient concentration binding experiment, four appropriate gradient concentrations of Rg3 (2.44, 7.3, 22, 66 µM) were selected, and the proper concentrations of the GRB2 proteins were all set at 1 mg/ml for the loading solution. The kinetic parameters, including *K_ON_*, *K_Off_* and *K_D_* were calculated by data analysis. The interaction was further validated with Microscale Thermophoresis (MST) methods using Monolith NT.115 (Nanotemper). Rg3 titrated in different concentrations to purified recombinant human GRB2 protein. The reaction was performed in 50 mM Hepes, 50 mM NaCl, 0.01% Tween 20 and 2 mM MgCl_2_. Then the samples were incubated in room temperature for 5 min before analyzing by microscale thermophoresis. In this instrument, an infra blue Laser (IB Laser) beam couples into the path of light (i.e. fluorescence excitation and emission) with a dichroic mirror and is focused into the sample fluid through the same optical element used for fluorescence imaging. The IB laser is absorbed by the aqueous solution in the capillary and locally heats the sample with a 1/e 2 diameter of 25 μm. Up to 24 mW of laser power where used to heat the sample, without damaging the biomolecules. To analyze the thermophoresis of a sample, ten microliters were transferred in a glass capillary (NanoTemper, hydrophilic treated). Thermophoresis of the protein in presence of varying concentrations of compound was analyzed for 30 seconds. Measurements were performed at room temperature and standard deviation was calculated from three independent experiments.

### Thermal shift assay

For the temperature-dependent thermal shift assay [7], 150 μL of lysates (3 mg/mL) from SHSY5Y cells were incubated with 5 μM of Rg3 at each temperature point from 43 to 60 °C for 3 min. Samples were repeatedly freezing and thawing by liquid nitrogen. Then, the samples were centrifuged at 20,000 g for 10 min at 4 °C to separate the supernatant and pellet. The supernatant was mixed with 5× loading buffer and then separated on a 10 % SDS-PAGE for immunoblotting analysis of GRB2.

### Prediction of transcription factor

To investigate the regulation of genes, we predicted the transcription factors (TFs) regulating cardiolipin synthase 1. The promoter sequences (upstream 2000 bp) of genes were downloaded from UCSC (http://genome.ucsc.edu/). Then we predicted the TFs and related TF binding sites (TFBS) using match tool from TRANSFAC (http://gene-regulation.com/pub/databases.html), which provides data on eukaryotic transcription factors and their experimentally-proven binding sites (table S7).

### Electrophoretic mobility shift assay (EMSA)

The probe set were designed and labeled with biotin to target the EVI1 binding sites on the CRLS1 promoter (table S8). Nuclear extracts from SH-SY5Y cells were incubated with biotin-labeled probes. A competition assay was conducted with either an unlabeled biotin probe containing a known EVI1 binding site (positive control) or an unlabeled probe with mutated EVI1 binding site. A supershift assay was performed with an EVI1 antibody.

### Chromatin immunoprecipitation (ChIP) and sequential ChIP assay

ChIP was performed with an EZ-ChIP kit as per the manufacturer’s instructions (table S9, Primers used in ChIP assays). Briefly, cells with different treatments were lysed in lysis buffer and sonicated (15 s on and 90 s off, repeated 8 times). After precipitation with Agarose A for 30 min, the fragmented DNA was pulled down with EVI1 antibody and then subjected to amplification by qPCR. Afterwards, sequential ChIP assays were performed. Briefly, after elution from protein A Sepharose beads, the DNA-protein complex was subjected to a second immunoprecipitation with EVI1 antibody. Control cell or rabbit IgG was included. GAPDH in DNA prior to immunoprecipitation was amplified as a control.

### Animals and experimental design

All experiments and animal care in this study were conducted in accordance with the National Institutes of Health guide for the care and use of Laboratory animals (NIH Publications No. 8023, revised 1978), and the Provision and General Recommendation of Chinese Experimental Animals Administration Legislation and approved by the Science and Technology Department of Jiangsu Province (SYXK (SU) 2016-0011). Male C57BL/6J mice (Yangzhou University, China) were housed on woodchip bedding in cages with 50–60% humidity and a 12 h light/dark cycle and given free access to food and water. In PD experiment, mice were randomly divided into 4 groups: control group, MPTP alone group, and two MPTP plus Rg3 treatment groups. The Grb2 knockdown fragment was ligated to an shRNA carrying AAV5 vector. The titers of adeno-associated virus (AAV) were between 1 × 10E13 and 2 × 10E13 vg/ml. Mice were anesthetized by intraperitoneal injection of 0.7% pentobarbital sodium and then fixed in a stereotaxic apparatus. A total of 2 × 10E9 units of scramble virus (control group, MPTP alone group, and one of MPTP plus Rg3 treatment groups) or virus targeting mouse Grb2 (another one of MPTP plus Rg3 treatment groups) in a 1 μl volume was slowly injected into the SN (AP: −3.3 mm; ML: ±1.3 mm; DV: −4.7 mm from bregma) at a rate of 0.2 μl/min using a 25 μl syringe. Control group mice were administered phosphate buffered saline (PBS) (vehicle) for 7 consecutive days by i.p. injection, while mice of the MPTP alone and MPTP plus baicalein groups were injected i.p. with 20 mg/kg MPTP (every 2 h for a total of four doses in 1 d) for 7 consecutive days to induce PD. Mice in the two MPTP plus Rg3 groups received intragastric Rg3 at 20 mg/kg/d 1 hour before MPTP treatment for 7 days and continued administration for an additional 2 days (total of 9 days), while Mice of the control group and MPTP alone group received intragastric 0.5% CMC-Na for 9 days in the same way. All mice were killed after behavioral testing conducted the day for immunohistochemistry and western blotting.

### Open Field Test

Open field tests were conducted as described [8]. The apparatus was made of white plastic (50 × 50 cm). Mice were transferred to the experimental room 1 hour before testing for adaptation and then placed into the open field chamber for 3 minutes. The total distance travelled was recorded automatically during test.

### Pole Test

Pole tests were conducted as described [8]. Briefly, a 0.5-m long pole was placed vertically into the home cage, and mice were placed at the top. The time required to descend back into the cage was recorded. On the day before the test, each mouse received 3 consecutive training trials, each separated by 30-minute rest periods. On the test day, 3 test trials were conducted with 3-minute intervals, and the average time to descend back into the cage was recorded.

### Accelerating Rotarod Test

Accelerating rotarod tests were conducted as described [8]. Briefly, mice were placed on the accelerating rotarod, which accelerates from 0 to 40 rpm in 180 seconds and the latency to fall off was recorded. Mice were pretrained over 3 consecutive days at a 30-minute interval. On the test day, mice were evaluated on 3 trials conducted with 3-minute intervals.

### Histological analysis

The fresh brain tissue was fixed with fixed liquid for more than 24 hours. Remove the tissue from the fixed liquid and trim the target tissue with a scalpel in the ventilation cupboard, and put the trimmed tissue and the label in the dehydration box. Dehydration and wax leaching: put the dehydration box into the dehydrator in order to dehydrate with gradient alcohol. 75% alcohol 4 hours, 85% alcohol 2 hours, 90% alcohol 2 hours, 95% alcohol 1 hour, anhydrous ethanol I 30 min, anhydrous ethanol II 30 min, alcohol benzene 5∼10 min, xylene II 5∼10 min, 65 °C melting paraffin I 1h, 65 °C melting paraffin II 1h, 65 °C melting paraffin III 1 hour. The wax-soaked tissue is embedded in the embedding machine. First, put the melted wax into the embedding frame, and before the wax solidifies, remove the tissue from the dewatering box and put it into the embedding frame according to the requirements of the embedding surface and affix the corresponding label. Cool at −20 °C freezing table, and after the wax is solidified, the wax block is removed from the embedded frame and repaired. Place the trimmed wax block on a paraffin slicer with a thickness of 4 μ m. The tissue is flattened when the slice floats on the 40 °C warm water of the spreading machine, and the tissue is picked up by the glass slides and baked in the oven at 60 °C. After the water-baked dried wax is melted, it is taken out and stored at room temperature. For immunofluorescent-based double labeling, the sequential staining procedure was followed. Brain sections were stained for Th and Il-1beta and images were acquired using a light microscopy (Ts2R, Nikon, Japan).

### Statistical analysis

Results are shown as means ± standard deviation. Significant differences between the groups were evaluated using the Student’s *t*-test. When three or more means are compared for statistical significance, one or two-way ANOVA was conducted with treatment or phenotype as the independent factors. When two groups of measurements were examined for statistical significance, the two-sided Student’s *t*-test was conducted. The survival curves of C. elegans lifespan were analyzed and compared using log-rank (Mantel-Cox) test.

---

**figure S1.**
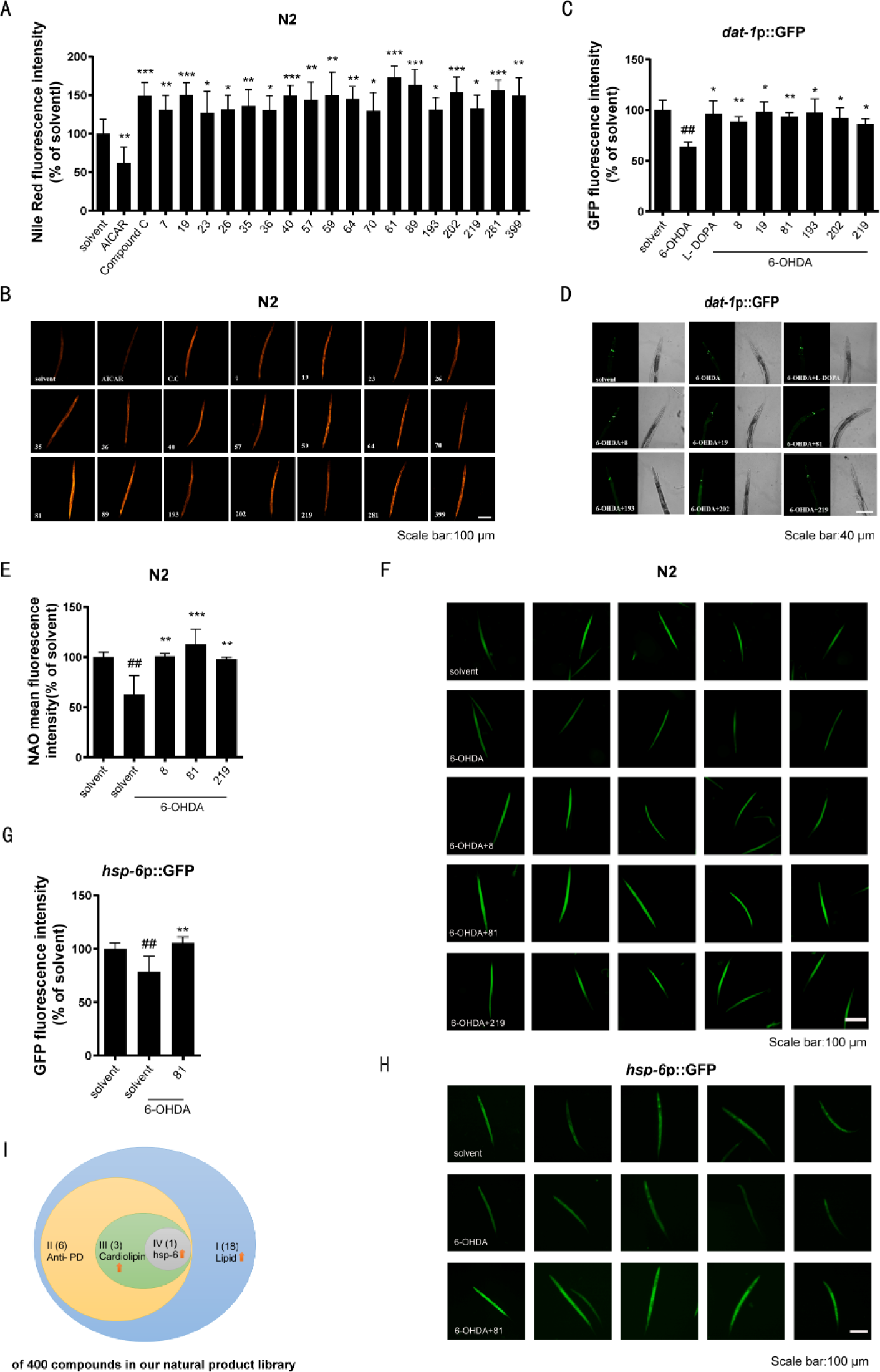
**(A-B)** The N2 strain was treated with corresponding natural products (10 µM) or AICAR (1 mM) or Compound C (20 µM) for 24 h. The lipid droplet was measured with Nile Red (1 μg/ml) staining. Scale bar: 100 µm. **(C-D)** The BZ555 strain was pretreated with 6-OHDA (30 mM) for 1 h, and then transferred to plates containing indicated compound (10 µM) and 5-Fluorouridine (0.04 mg/ml) for 72 h. The fluorescence intensity of dopaminergic neurons was measured. Scale bar: 40 µm. **(E-F)** The N2 strain was pretreated with 6-OHDA (30 mM) for 1 h, then the nematodes were transfered to the plates containing indicated compound (10 µM) for 36h, followed by staining with NAO (10 μM). The fluorescence intensity of cardiolipin was measured. Scale bar: 100 µm. **(G-H)** The SJ4100 strain was pretreated with 6-OHDA (30 mM) for 1 h, then the nematodes were transferred to the plates containing indicated compound (10 µM) for 12 h. The fluorescence intensity of nematodes was measured using inverted fluorescence microscopy. Scale bar: 100 µm. **(I)** Schematic diagram of screening results. ## *p* < 0.01 vs solvent-only control, \**p* < 0.05, \*\**p* < 0.01, \*\*\**p* < 0.001 vs 6-OHDA control (n=5-8).

**figure S2.**
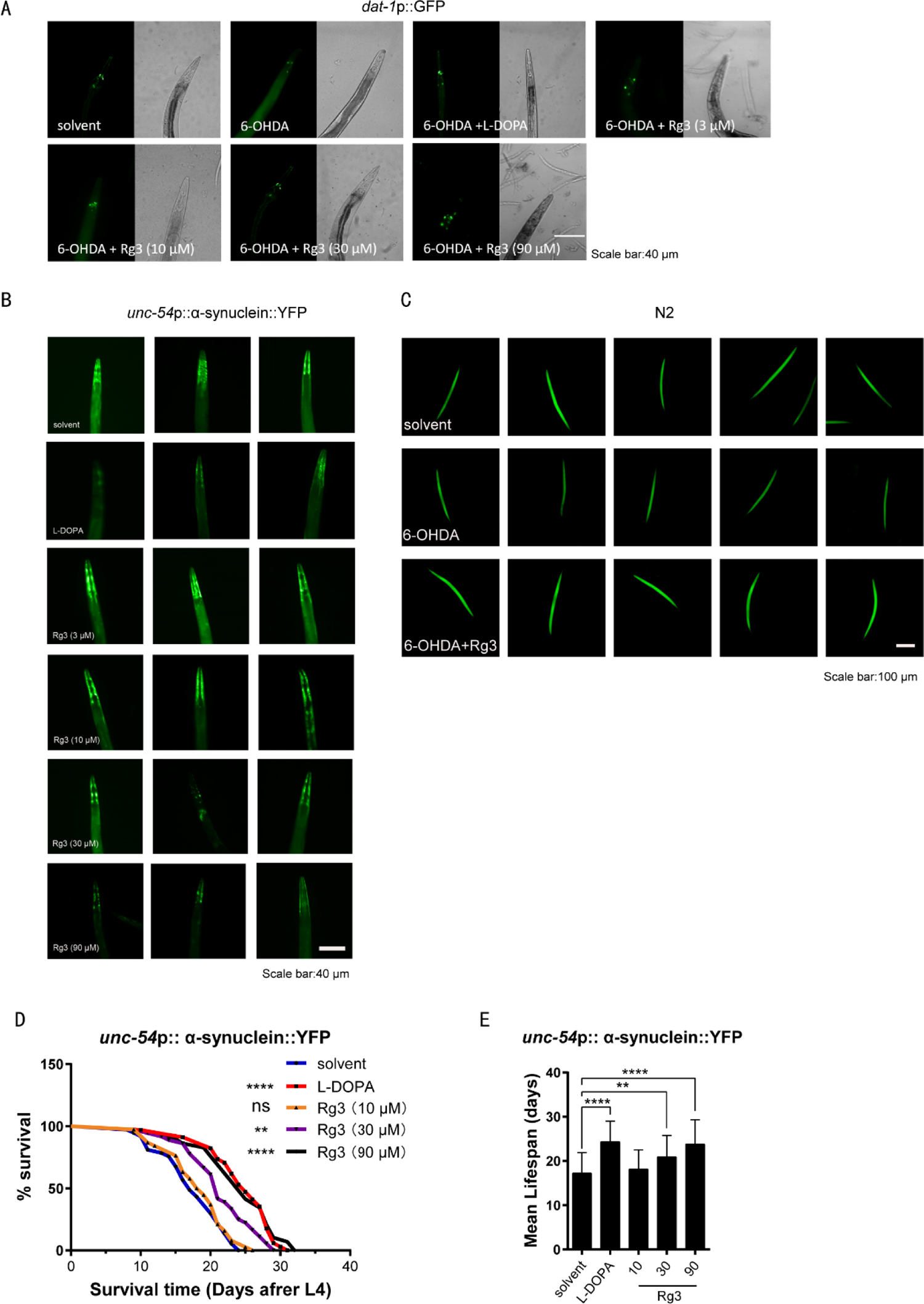
**(A)** The BZ555 strain was pretreated with 6-OHDA (30 mM) for 1 h then transferred to the plates with 5-Fluorouridine (0.04 mg/mL) and indicated concentrations of Rg3 or L-DOPA (10 µM) for 72 h. The fluorescence intensity of dopaminergic neurons was measured. Scale bar: 40 µm. **(B)** The eggs of NL5901 strain were treated with indicated concentrations of Rg3 or L-DOPA (10 µM). After growing to the L4 stage, the nematodes were treated with 5-Fluorouridine (0.04 mg/mL) for 12 days. The fluorescence intensity of α-synuclein protein in nematodes was measured. Scale bar: 40 µm. **(C)** The N2 strain was pretreated 6-OHDA (30 mM) for 1 h then transferred to the plates with Rg3 (90 µM) for 36h. The cardiolipin content was measured by NAO (10 μM) fluorescence signaling. Scale bar: 100 µm. **(D-E)** The eggs of NL5901 strain were treated with indicated concentrations of Rg3 or L-DOPA (10 µM). After growing to the L4 stage, the nematodes were treated with 5-Fluorouridine (0.04 mg/mL). After 60 h, the number of nematodes was recorded every day. \*\**p* < 0.01, \*\*\**p* < 0.001, \*\*\*\**p* < 0.0001 vs. indicated control (A-C: n=5-12; D-E: n=29-38).

**figure S3.**
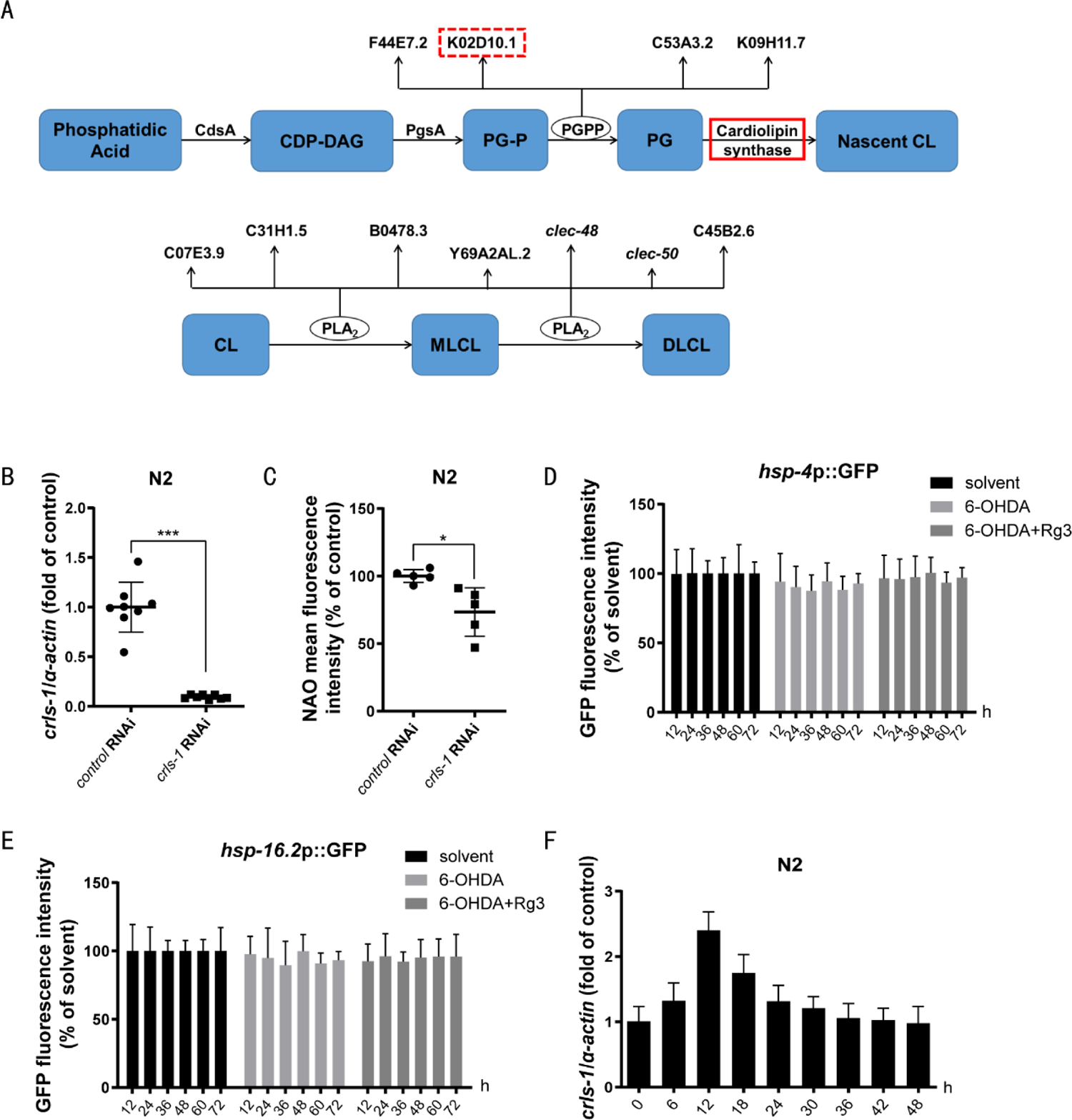
**(A)** Genes involved in cardiolipin synthesis and hydrolysis. **(B)** The mRNA levels of *crls1* were measured in the N2 strain treated with *crls1* RNAi. **(C)** The *crls1* RNAi-treated N2 strain was stained with NAO (10 µM). Cardiolipin content was measured by the NAO fluorescence intensity. **(D-E)** The SJ4005 **(D)** and CL2070 strains **(E)** were treated with 6-OHDA (30 mM) for different durations. The green fluorescence signals representing UPR^ER^ and HSR were measured respectively. **(F)** The N2 strain was treated with Rg3 (90 µM) for different durations. The mRNA levels of *crls1* were measured by qRT-PCR. \**p* < 0.05, \*\*\**p* < 0.001 vs. indicated control (n=4-8).

**figure S4.**
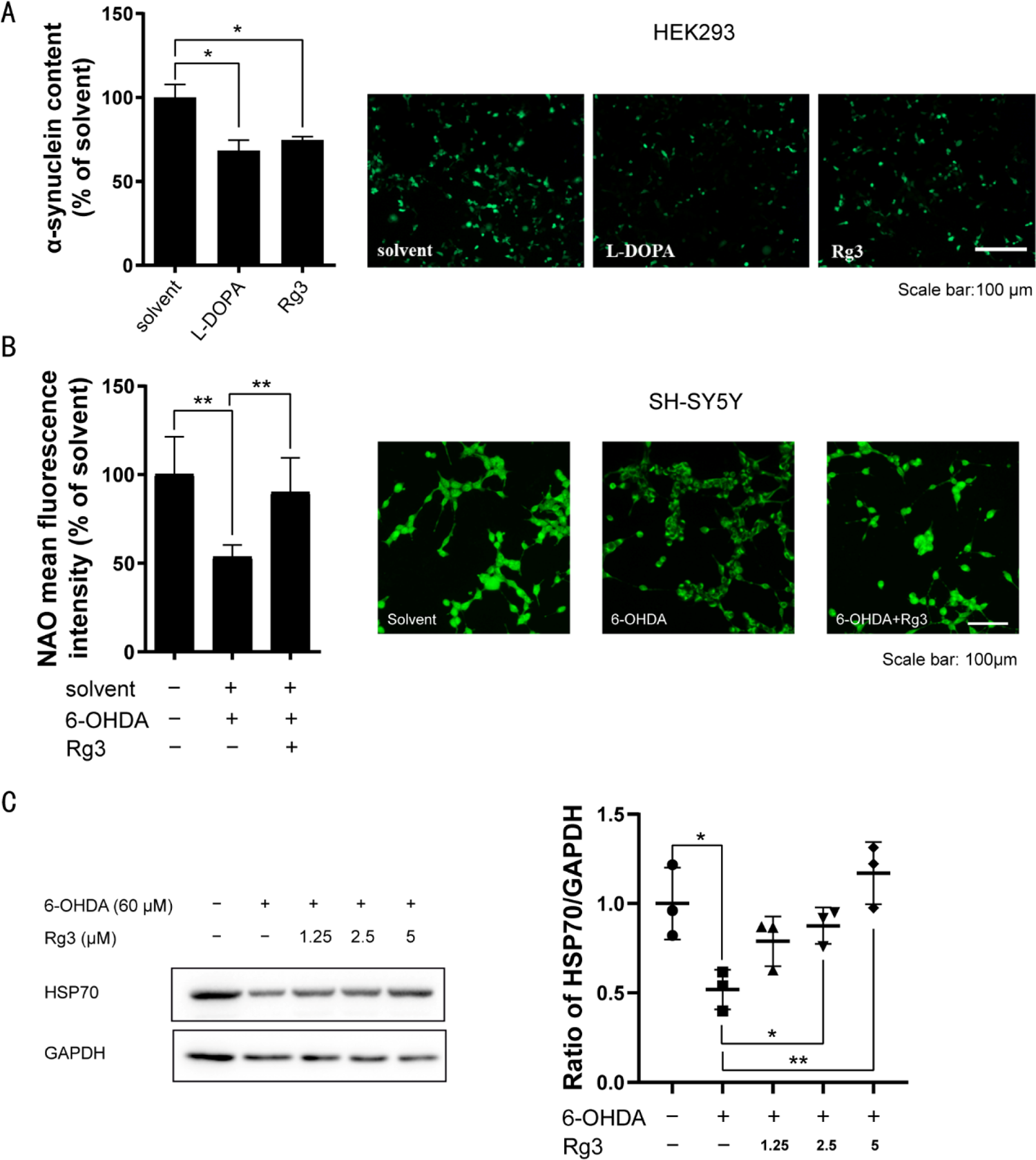
**(A)** HEK293 cells were transfected with GFP-tagged α-synuclein plasmid, and then treated with Rg3 (5 μM) or L-DOPA (10 μM) for 48 h. The amount of α-synuclein protein level was measured by green fluorescence signals. Scale bar: 100 µm. **(B-C)** SH-SY5Y cells were treated with Rg3 (5 μM) and 6-OHDA (60 μM) for 24 h: **(B)** Fluorescence intensity of cells was measured by NAO (5 μM) fluorescence signaling. Scale bar: 100 µm. **(C)** Total proteins were extracted and subjected to western blot with indicated antibodies. All experiments were repeated at least three times, \**p* < 0.05, \*\**p* < 0.01 vs. indicated control.

**figure S5.**
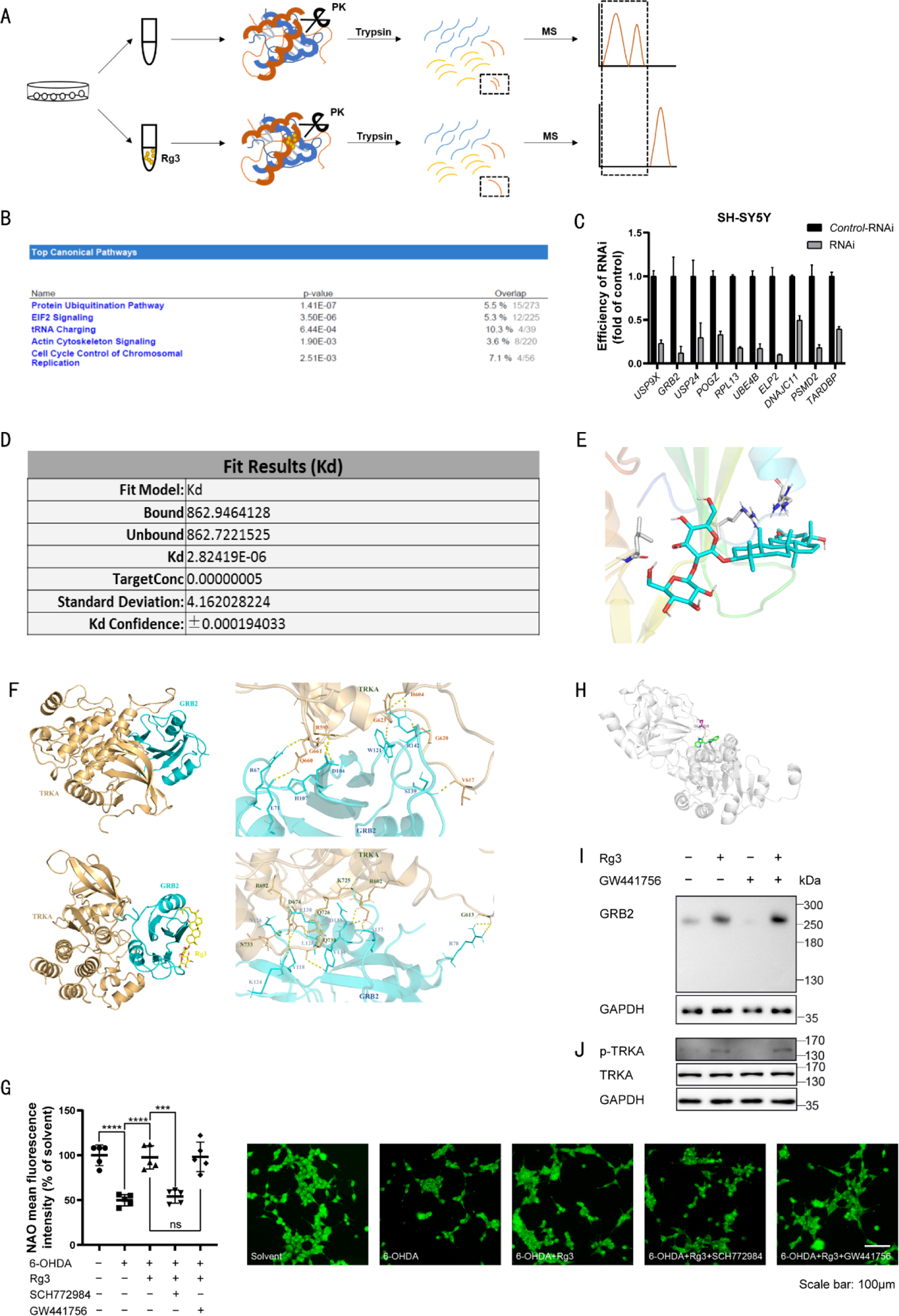
**(A)** Diagram of LiP-SMap strategy. **(B)** Ingenuity pathway analysis of the 253 proteins. **(C)** SH-SY5Y cells were transfected with indicated siRNA for 6 h, then cultured with normal medium for 48h. The mRNA levels of indicated targets were measured by qRT-PCR. **(D)** Measurement of *K_D_* value of the direct binding of GRB2 protein and Rg3 by Microscale Thermophoresis (MST). **(E-F)** *In silico* molecular docking using Discovery Studio**. (E)** *In silico* molecular docking of the direct binding of Rg3 and GRB2 protein. **(F)** *In silico* molecular docking of the direct binding of TRKA and GRB2 protein (top). *In silico* molecular docking of the direct binding of TRKA and GRB2-Rg3 complex (below). **(G)** SH-SY5Y cells were treated with 6-OHDA (60 μM) and 0.1% DMSO, Rg3 (5 μM), Rg3 (5 μM) + SCH772984 (10 μM) or Rg3 (5 μM) + GW441756 (10 μM) as indicated for 24 h. The cardiolipin content was measured by NAO (5 μM) fluorescence signaling. Scale bar: 100 µm. **(H)** *In silico* molecular docking of the direct binding of TRKA protein and GW441756 (Autodock 4.2). **(I-J)** SH-SY5Y cells were treated with Rg3 (5 μM) and TRKA inhibitor GW441756 (10 μM) as indicated for 3 h. **(I)** Then protein samples of each group were collected and applied for Native-PAGE (GRB2) and WB (GAPDH) analysis. **(J)** Total proteins were extracted and subjected to western blot with indicated antibodies. \*\*\**p* < 0.001, \*\*\*\**p* < 0.0001 vs. indicated control.

**figure S6.**
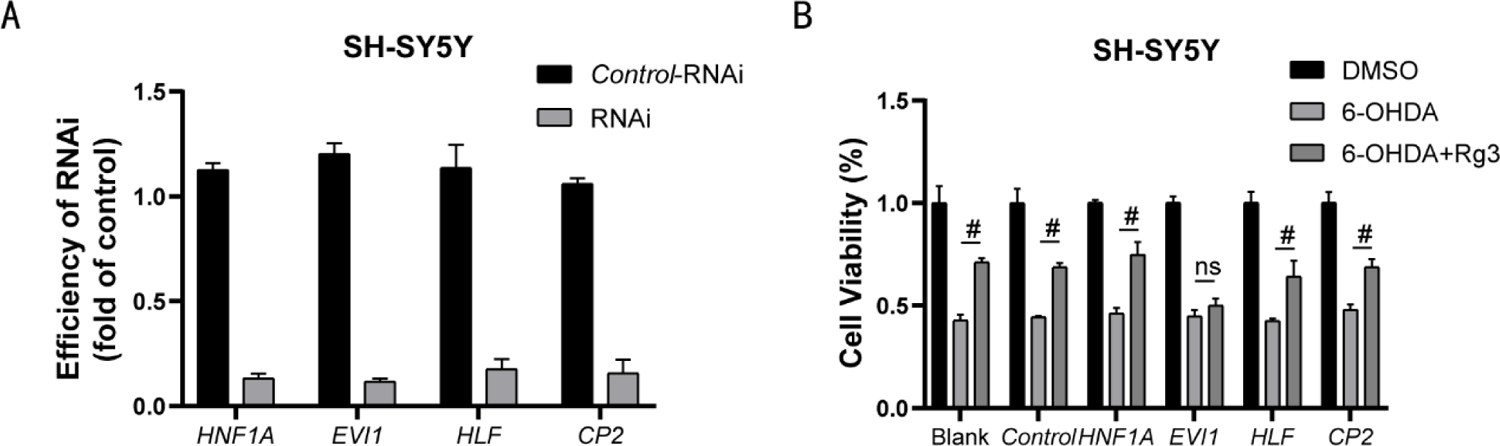
**(A)** SH-SY5Y cells were transfected with indicated siRNAs for 6 h, then cultured with normal medium for 48h. The mRNA levels of indicated targets were measured. **(B)** After transfected with indicated siRNAs, cells were treated with Rg3 (5 μM) and 6-OHDA (60μM) as indicated for 24 h. Cell viability was measured by CCK8 assay. All experiments were repeated at least three times. #*p* < 0.0001 vs. indicated control.

**figure S7.**
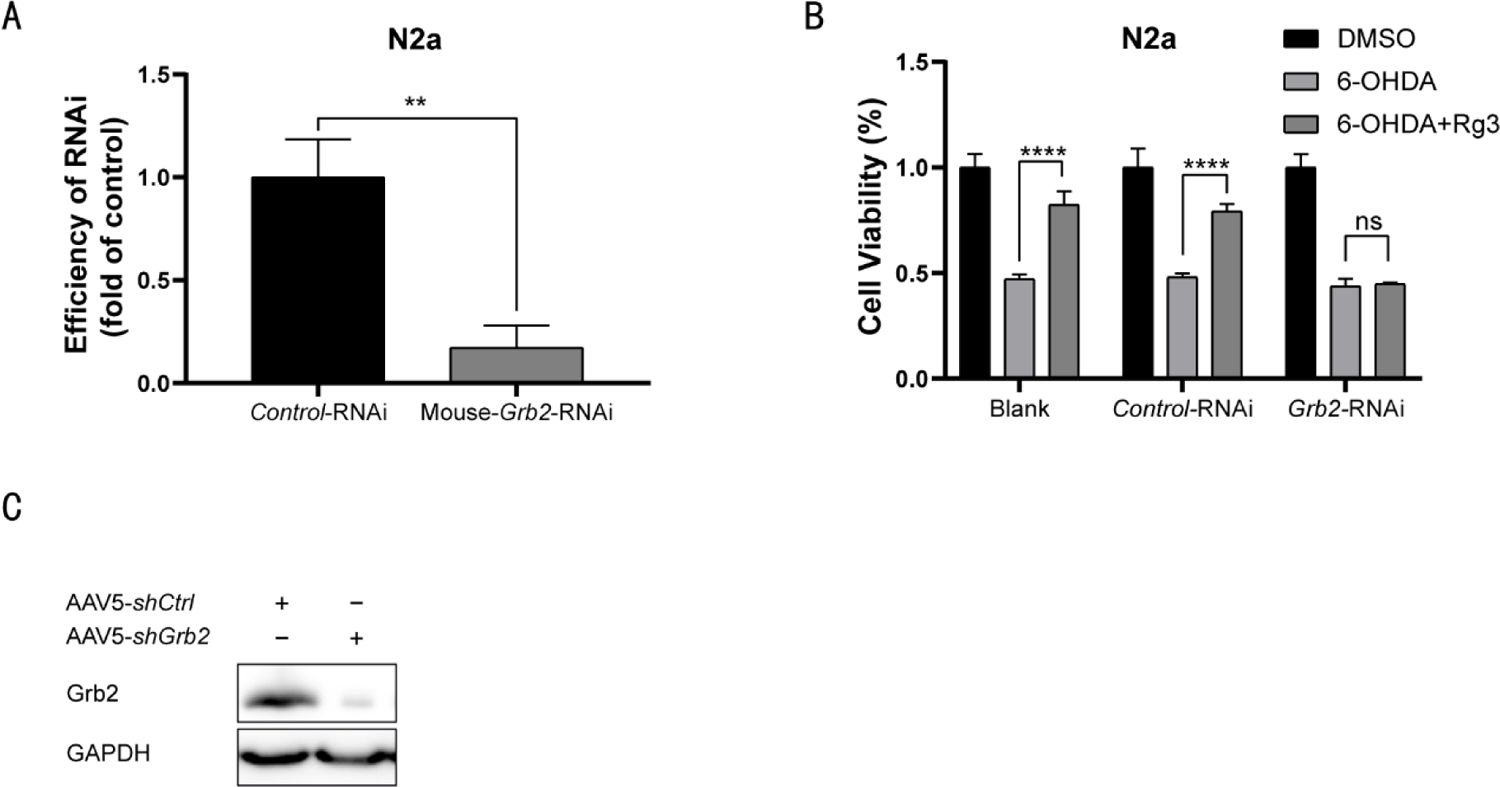
**(A)** N2a cells were transfected with indicated siRNAs for 6 h, then cultured with normal medium for 48h. The mRNA levels of indicated targets were measured. **(B)** After transfected with indicated siRNAs, cells were treated with Rg3 (5 μM) and 6-OHDA (60μM) as indicated for 24 h. Cell viability was measured by CCK8 assay. **(C)** Immunoblotting analysis of Grb2 and Gapdh in the SN from 4 groups of mice as indicated. All experiments were repeated at least three times. **p < 0.01, ****p < 0.0001 vs. indicated control.

**figure S8.**
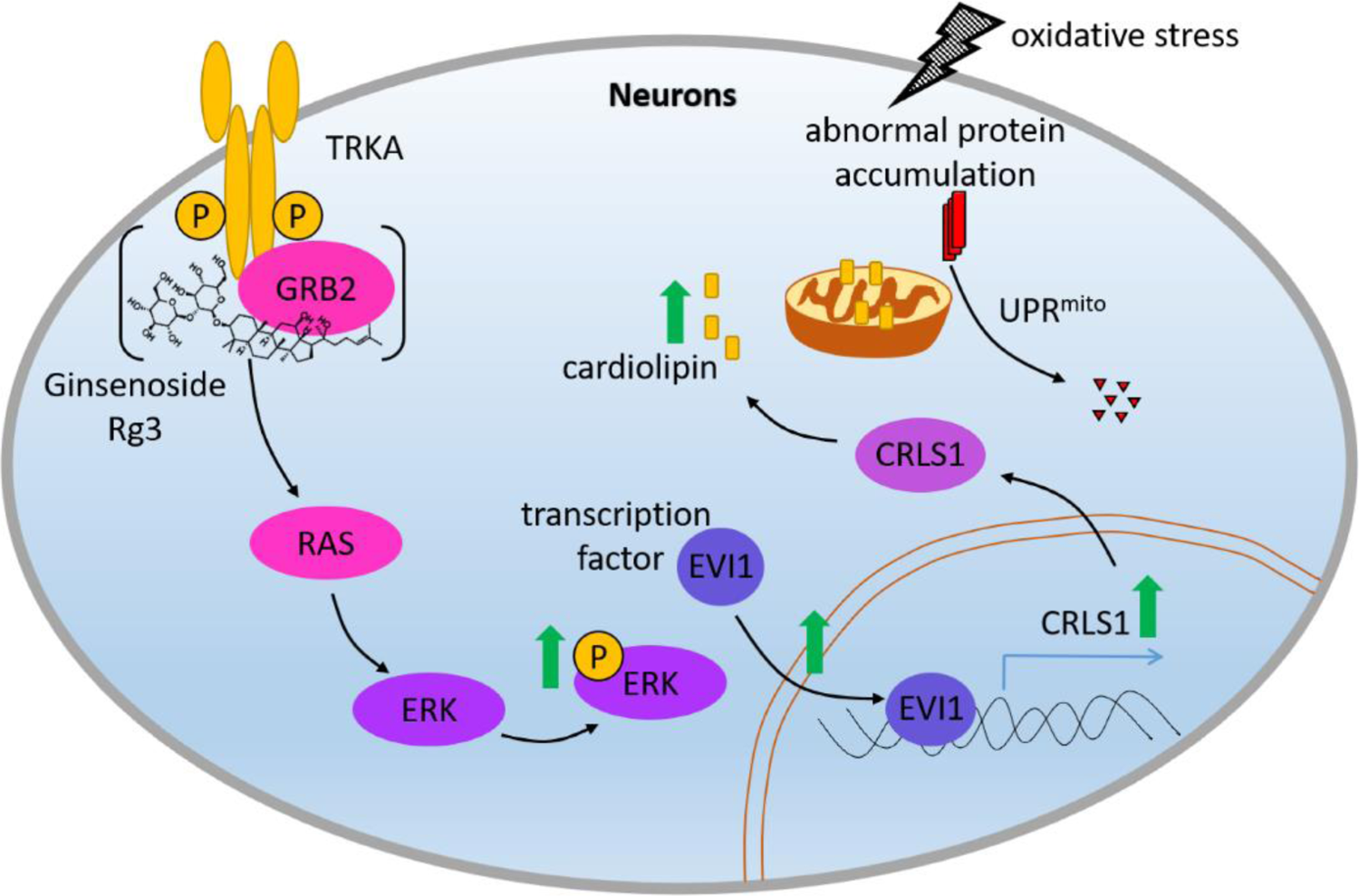
Proposed mechanism of Rg3-mediated neuroprotective effects.

**table S1.**
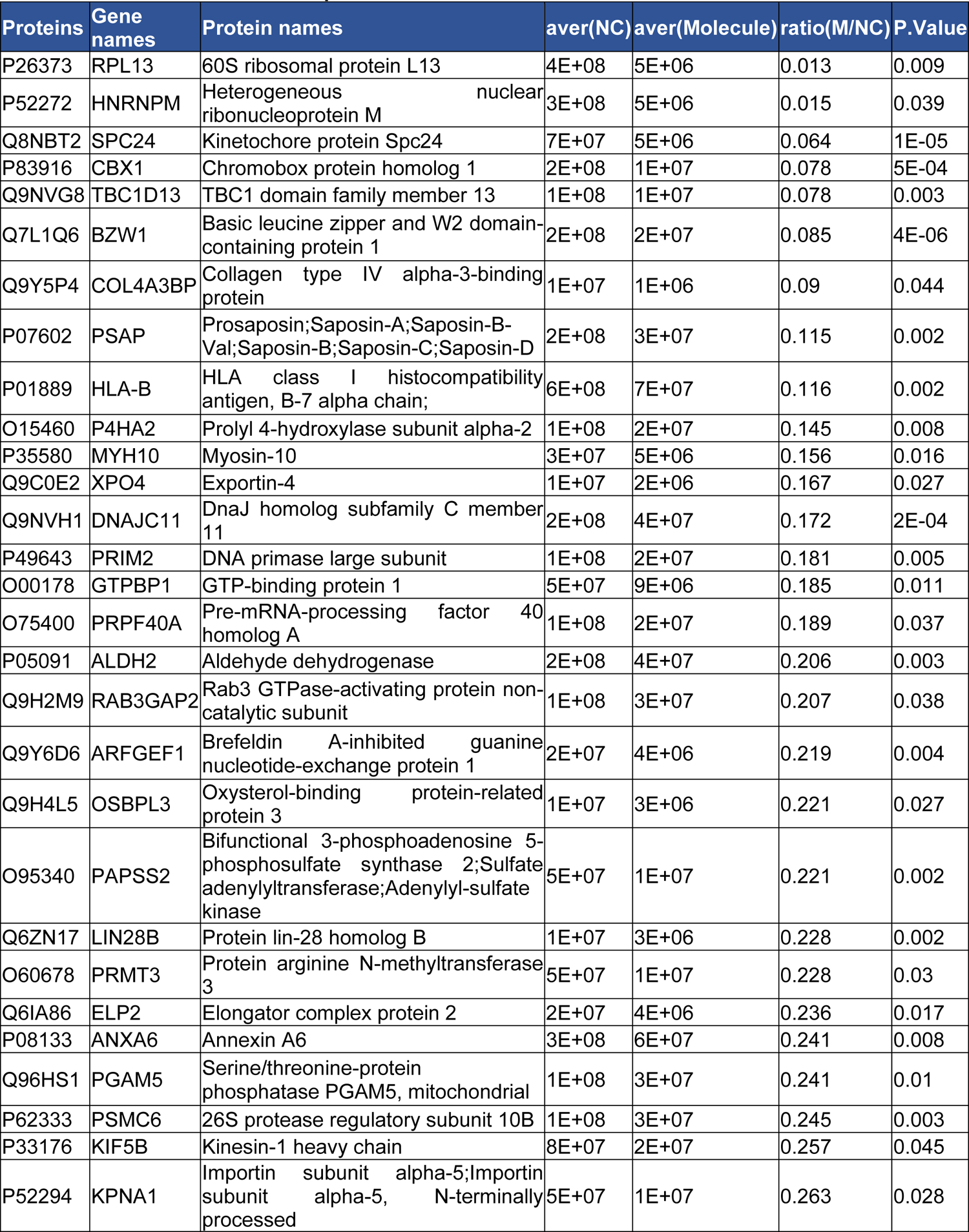
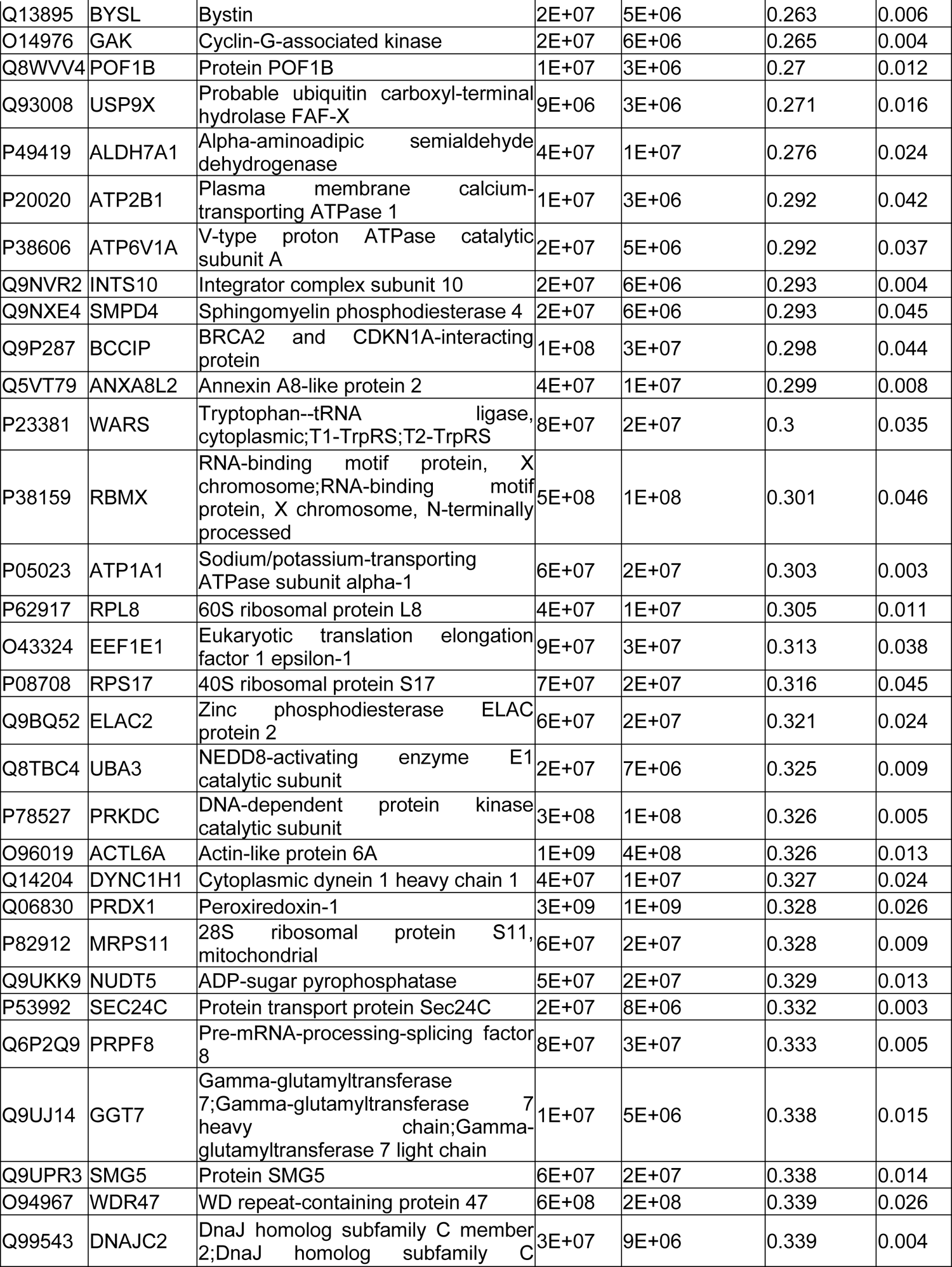
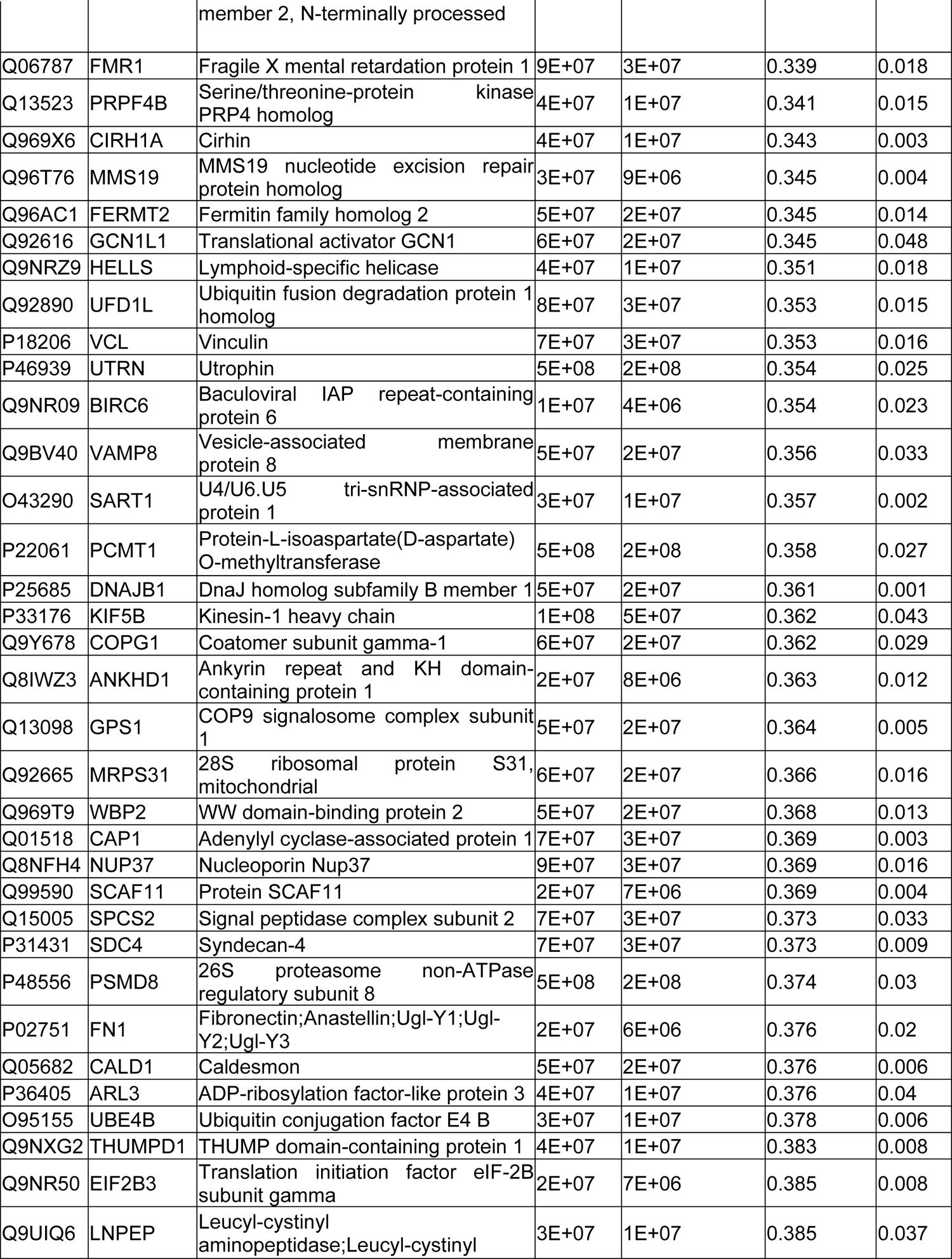
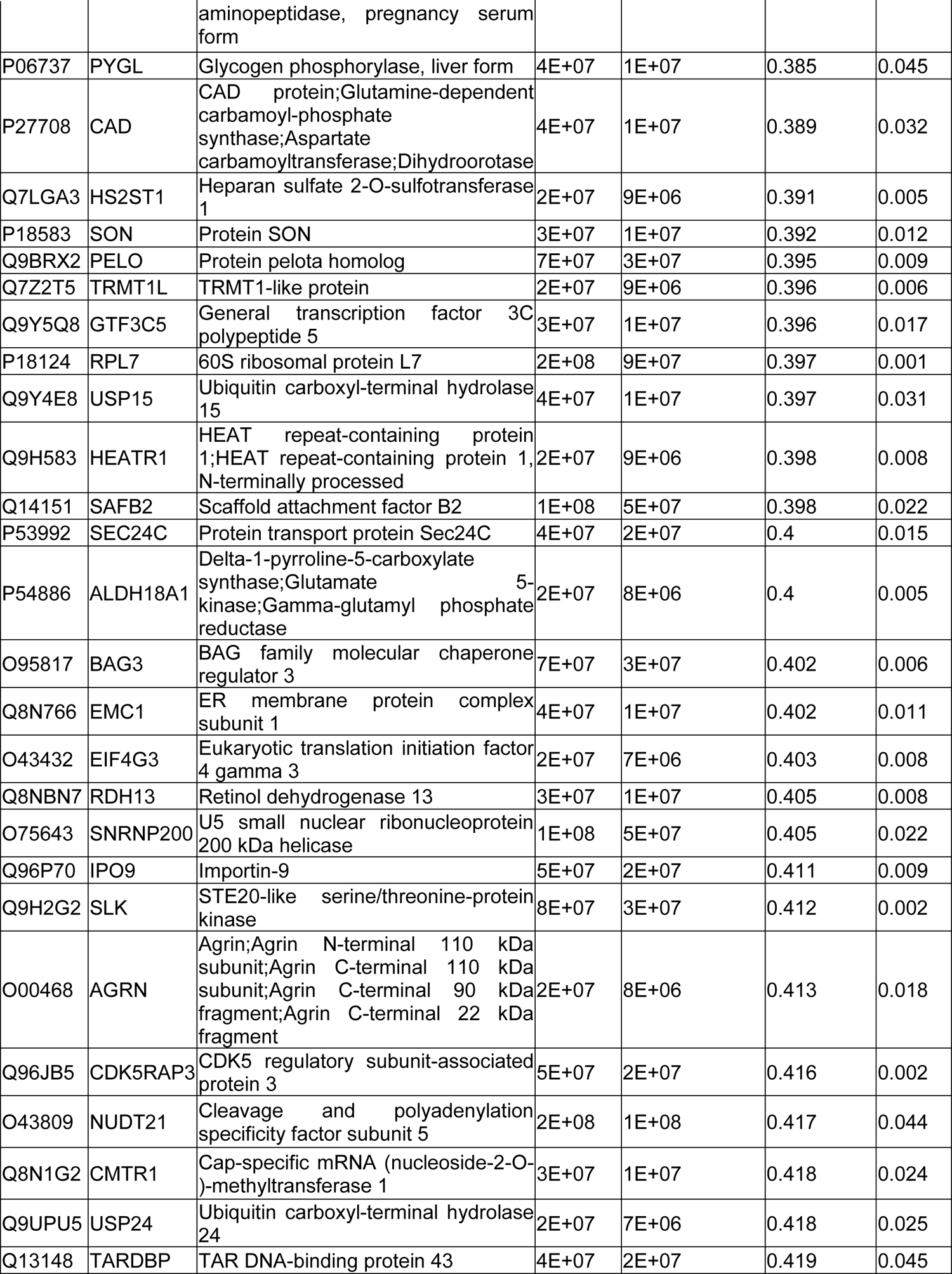
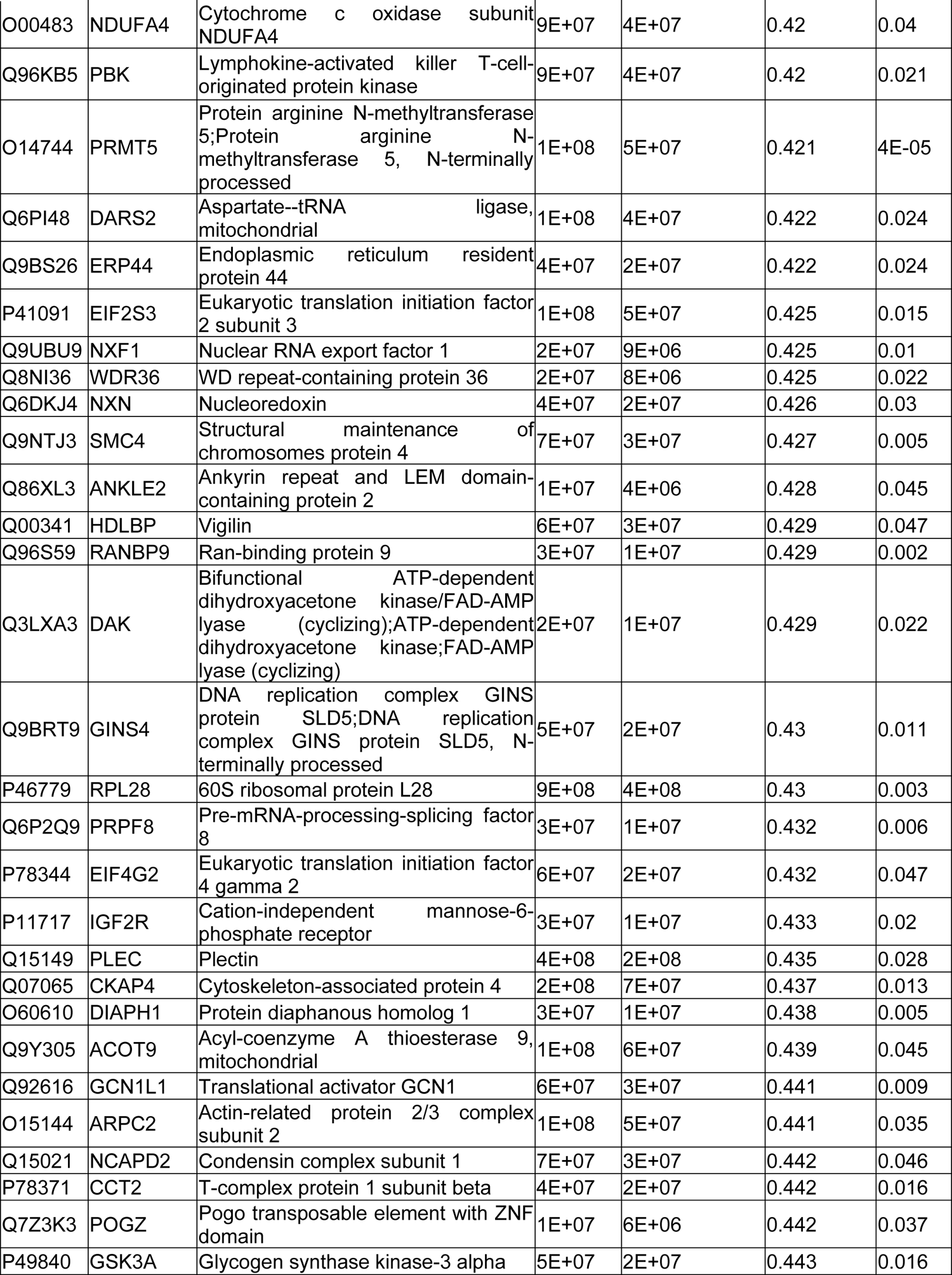
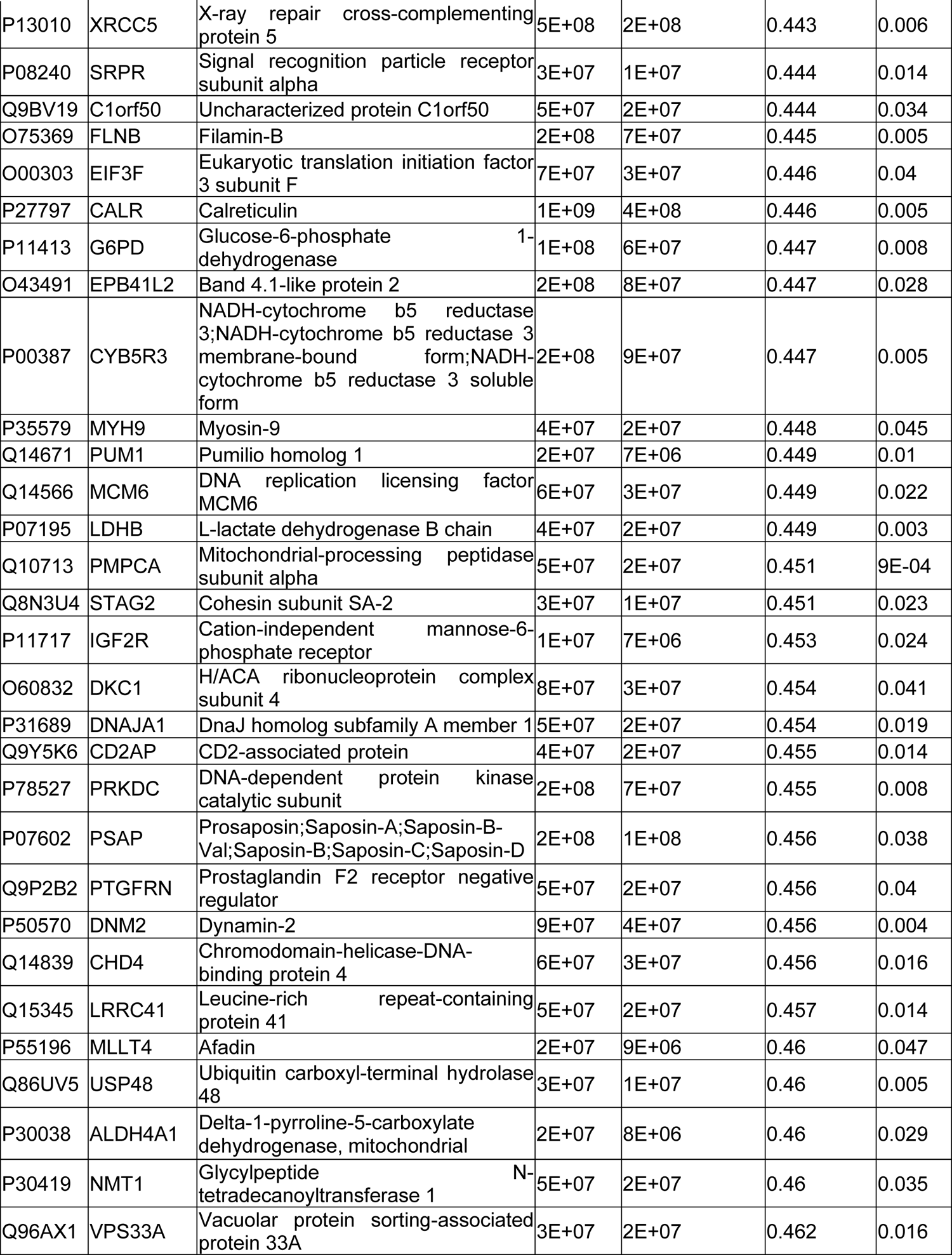
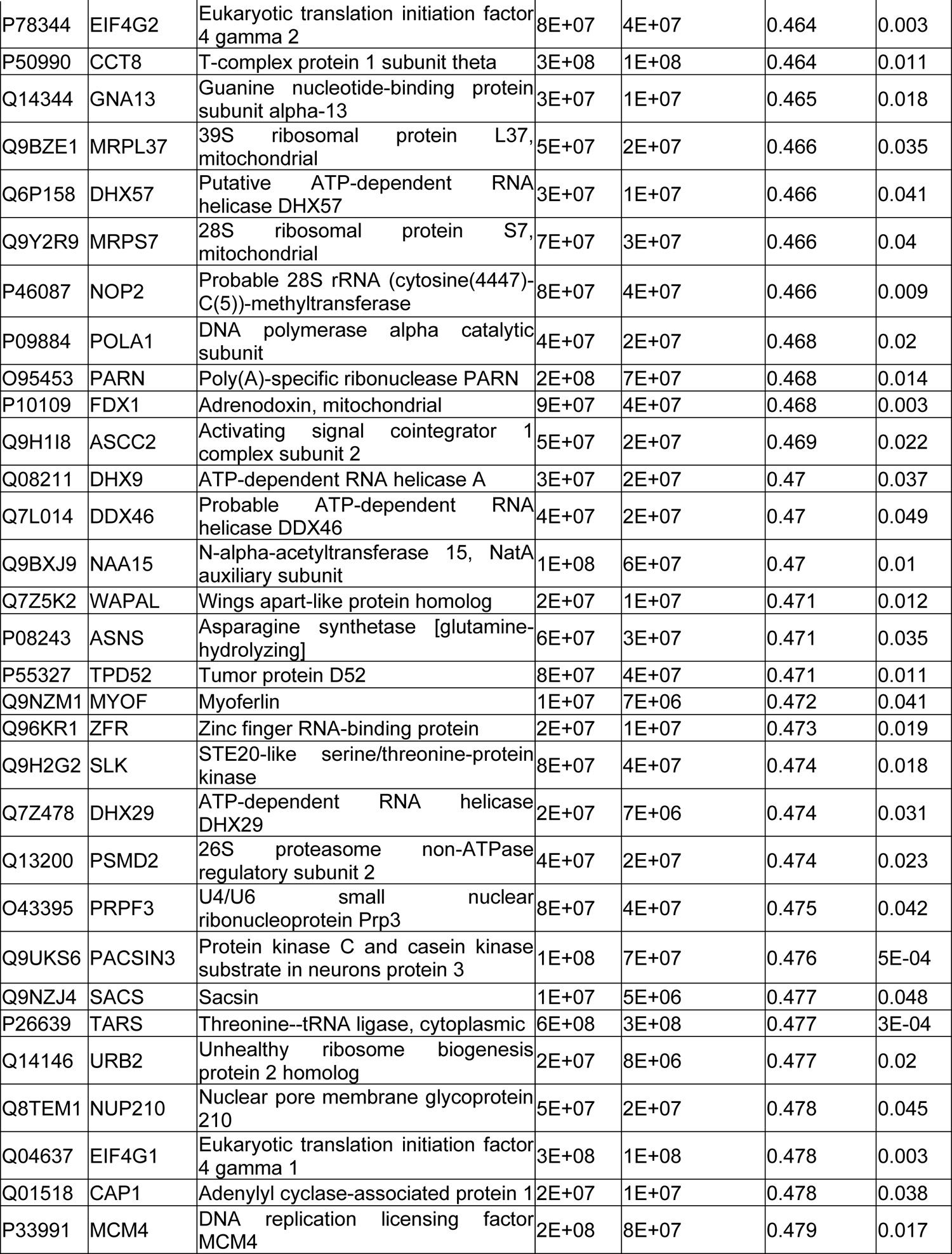
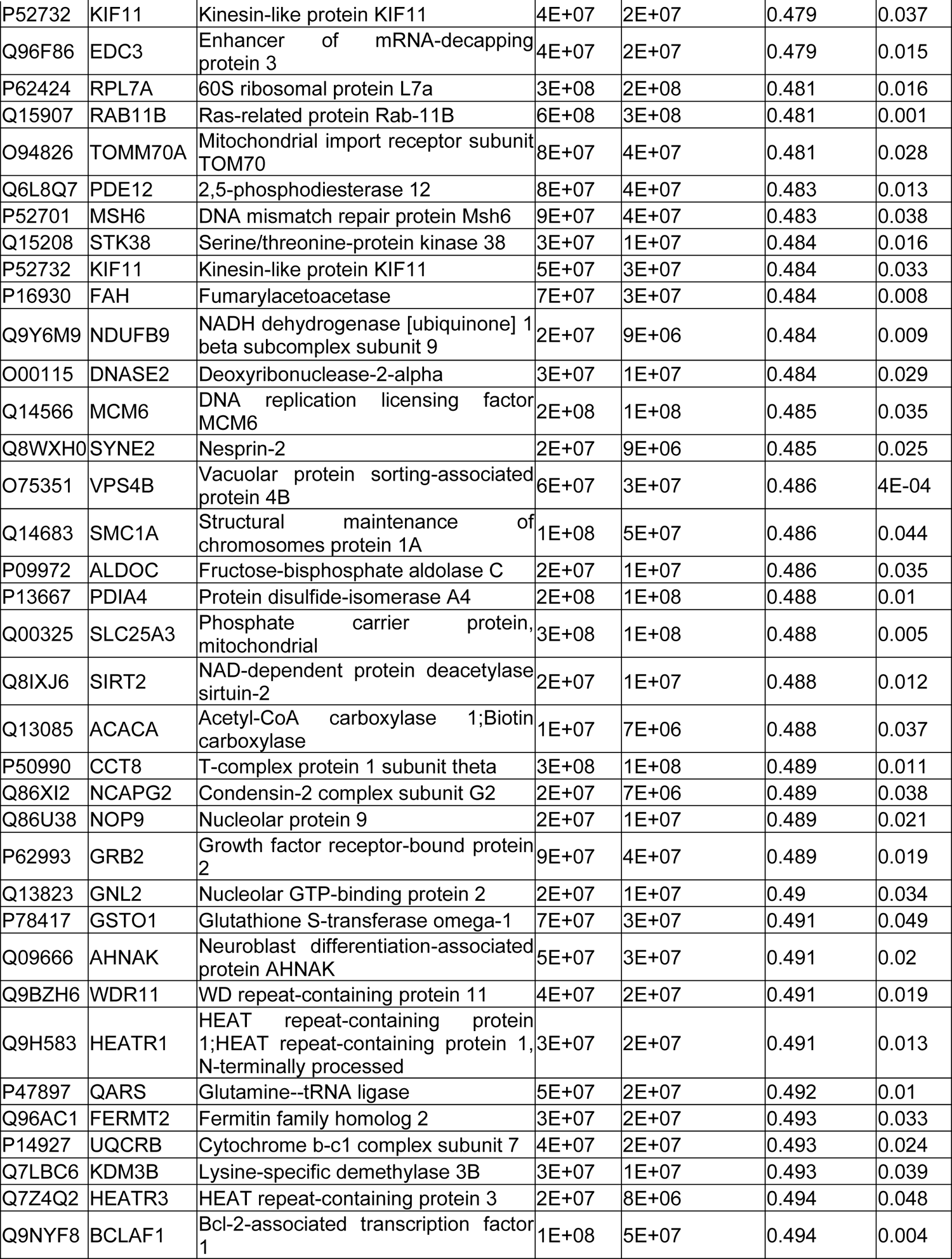
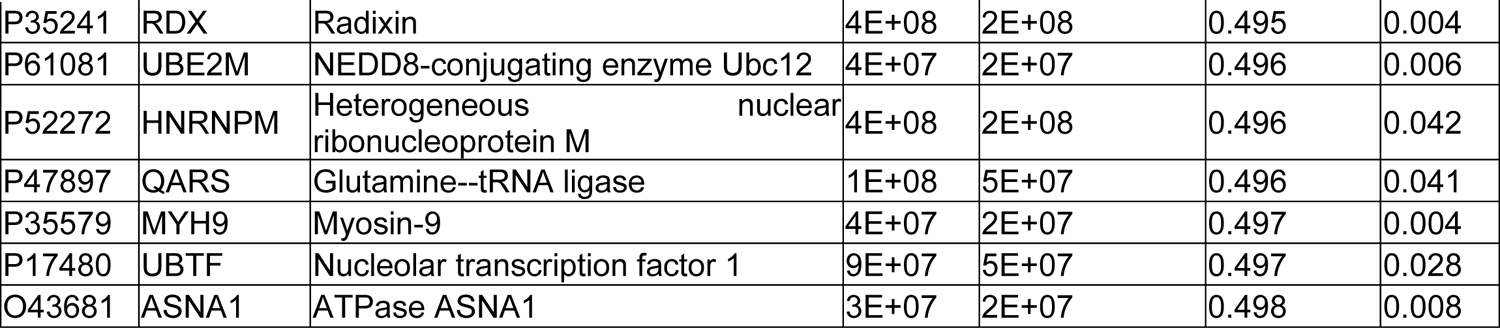
The result of LipSMAP

**table S2.** List of the candidate genes that related to neurodegenerative diseases

| NO. | Gene names | Official Full Name<br>(provided by HGNC) |
| --- | --- | --- |
| 1 | *ELP2* | elongator acetyltransferase complex subunit 2 |
| 2 | *POGZ* | pogo transposable element derived with ZNF domain |
| 3 | *TARDBP* | TAR DNA binding protein |
| 4 | *USP9X* | ubiquitin specific peptidase 9 X-linked |
| 5 | *DNAJC11* | DnaJ heat shock protein family (Hsp40) member C11 |
| 6 | *PSMD2* | proteasome 26S subunit, non-ATPase 2 |
| 7 | *UBE4B* | ubiquitination factor E4B |
| 8 | *USP24* | ubiquitin specific peptidase 24 |
| 9 | *GRB2* | growth factor receptor bound protein 2 |
| 10 | *RPL13* | ribosomal protein L13 |

**table S3.**
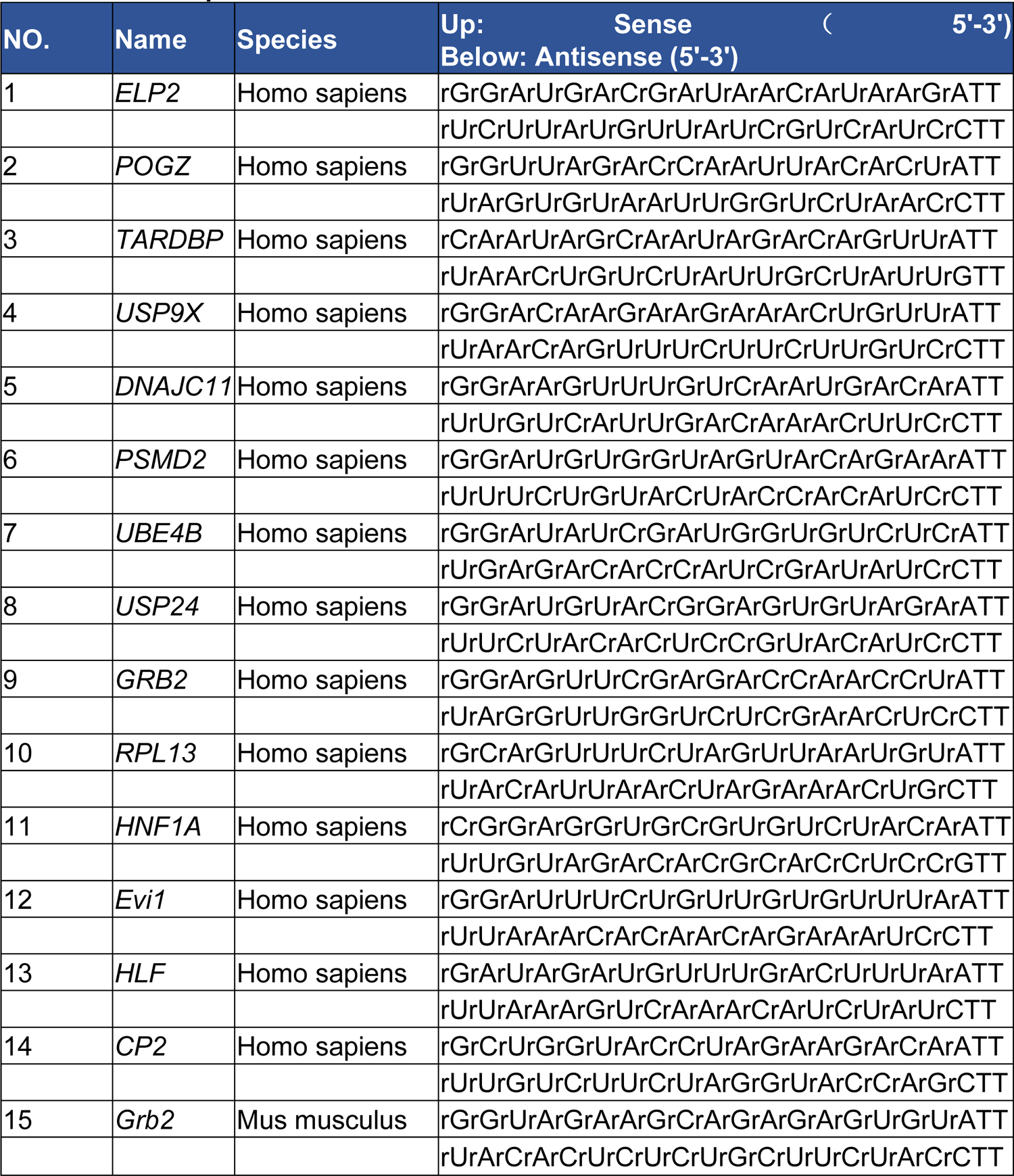
The sequence of siRNAs used

**table S4.**
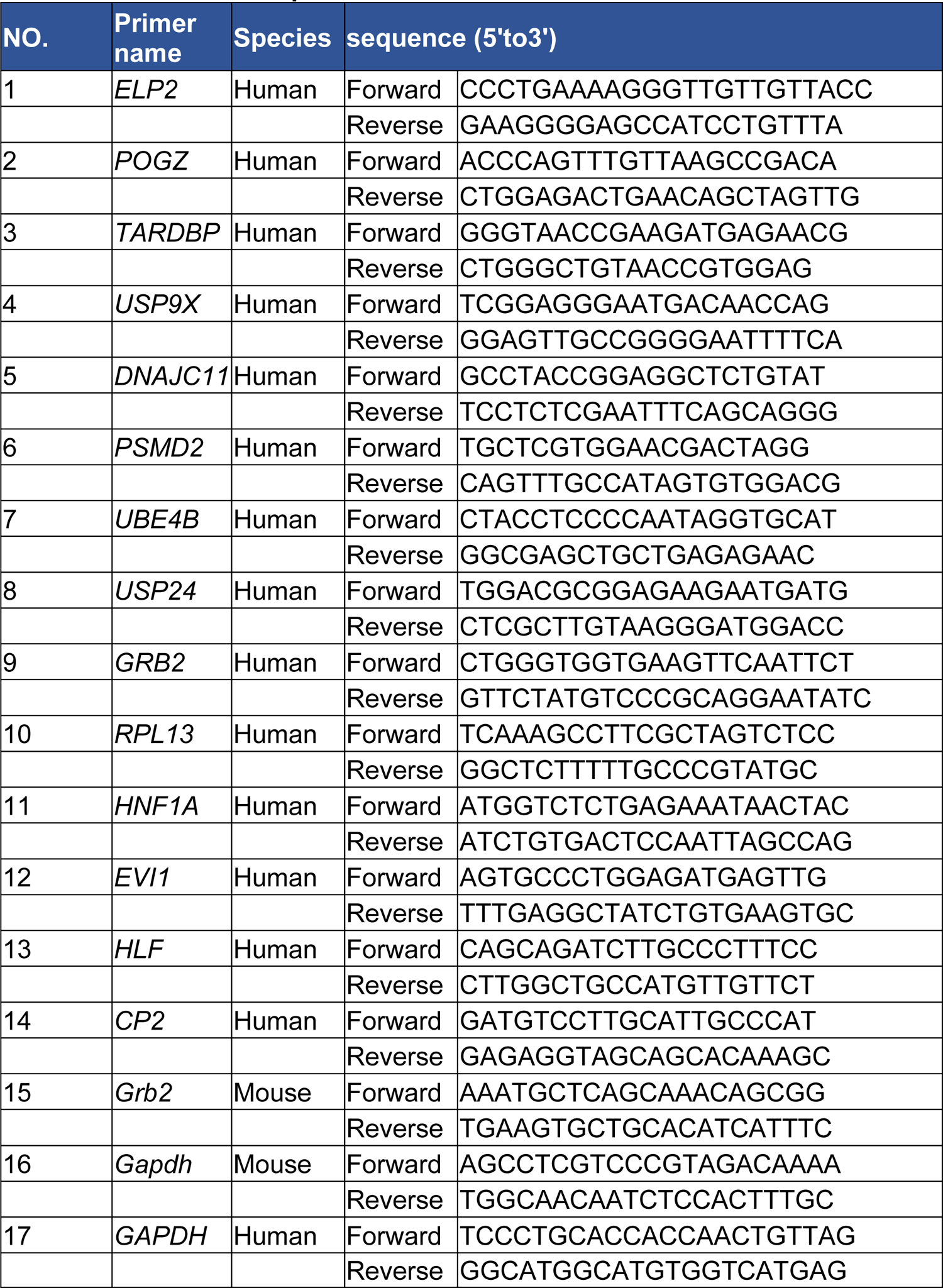
Primers used in qRT-PCR

**table S5.** The hydrogen bonding interaction between GRB2 and TRKA

| Receptor Residue | Ligand Residue |  | Distance | Type |
| --- | --- | --- | --- | --- |
| 4AOJ:ARG593 | 1CJ1:ASP104 | 4AOJ:ARG593:HH21 - 1CJ1:ASP104:OD2 | 2.3826 | Conventional |
| 4AOJ:GLY623 | 1CJ1:TRP121 | 4AOJ:GLY623:HN - 1CJ1:TRP121:O | 2.6321 | Conventional |
| 4AOJ:GLN660 | 1CJ1:GLU71 | 4AOJ:GLN660:HE22 - 1CJ1:GLU71:OE2 | 2.1434 | Conventional |
| 4AOJ:GLY661 | 1CJ1:HIS107 | 4AOJ:GLY661:HN - 1CJ1:HIS107:O | 2.6537 | Conventional |
| 4AOJ:GLN660 | 1CJ1:ARG67 | 1CJ1:ARG67:HH22 - 4AOJ:GLN660:O | 2.1862 | Conventional |
| 4AOJ:VAL617 | 1CJ1:SER139 | 1CJ1:SER139:HN - 4AOJ:VAL617:O | 2.0272 | Conventional |
| 4AOJ:GLY620 | 1CJ1:ARG142 | 1CJ1:ARG142:HH11 - 4AOJ:GLY620:O | 1.9331 | Conventional |
| 4AOJ:HIS604 | 1CJ1:ARG142 | 1CJ1:ARG142:HH12 - 4AOJ:HIS604:O | 2.0087 | Conventional |
| 4AOJ:HIS604 | 1CJ1:ARG142 | 1CJ1:ARG142:HH22 - 4AOJ:HIS604:O | 2.5049 | Conventional |

**table S6.**
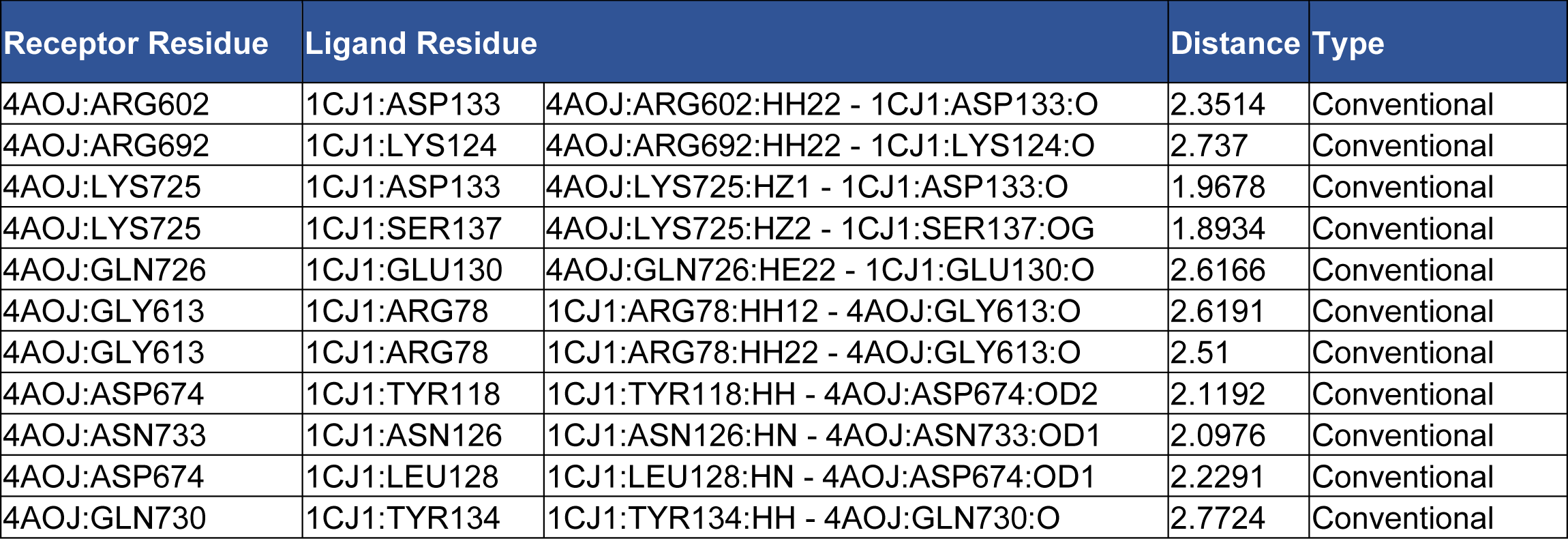
The hydrogen bonding interaction between TRKA and GRB2-Rg3 complex

| Receptor Residue | Ligand Residue |  | Distance | Type |
| --- | --- | --- | --- | --- |
| 4AOJ:ARG602 | 1CJ1:ASP133 | 4AOJ:ARG602:HH22 - 1CJ1:ASP133:O | 2.3514 | Conventional |
| 4AOJ:ARG692 | 1CJ1:LYS124 | 4AOJ:ARG692:HH22 - 1CJ1:LYS124:O | 2.737 | Conventional |
| 4AOJ:LYS725 | 1CJ1:ASP133 | 4AOJ:LYS725:HZ1 - 1CJ1:ASP133:O | 1.9678 | Conventional |
| 4AOJ:LYS725 | 1CJ1:SER137 | 4AOJ:LYS725:HZ2 - 1CJ1:SER137:OG | 1.8934 | Conventional |
| 4AOJ:GLN726 | 1CJ1:GLU130 | 4AOJ:GLN726:HE22 - 1CJ1:GLU130:O | 2.6166 | Conventional |
| 4AOJ:GLY613 | 1CJ1:ARG78 | 1CJ1:ARG78:HH12 - 4AOJ:GLY613:O | 2.6191 | Conventional |
| 4AOJ:GLY613 | 1CJ1:ARG78 | 1CJ1:ARG78:HH22 - 4AOJ:GLY613:O | 2.51 | Conventional |
| 4AOJ:ASP674 | 1CJ1:TYR118 | 1CJ1:TYR118:HH - 4AOJ:ASP674:OD2 | 2.1192 | Conventional |
| 4AOJ:ASN733 | 1CJ1:ASN126 | 1CJ1:ASN126:HN - 4AOJ:ASN733:OD1 | 2.0976 | Conventional |
| 4AOJ:ASP674 | 1CJ1:LEU128 | 1CJ1:LEU128:HN - 4AOJ:ASP674:OD1 | 2.2291 | Conventional |
| 4AOJ:GLN730 | 1CJ1:TYR134 | 1CJ1:TYR134:HH - 4AOJ:GLN730:O | 2.7724 | Conventional |

**table S7.**
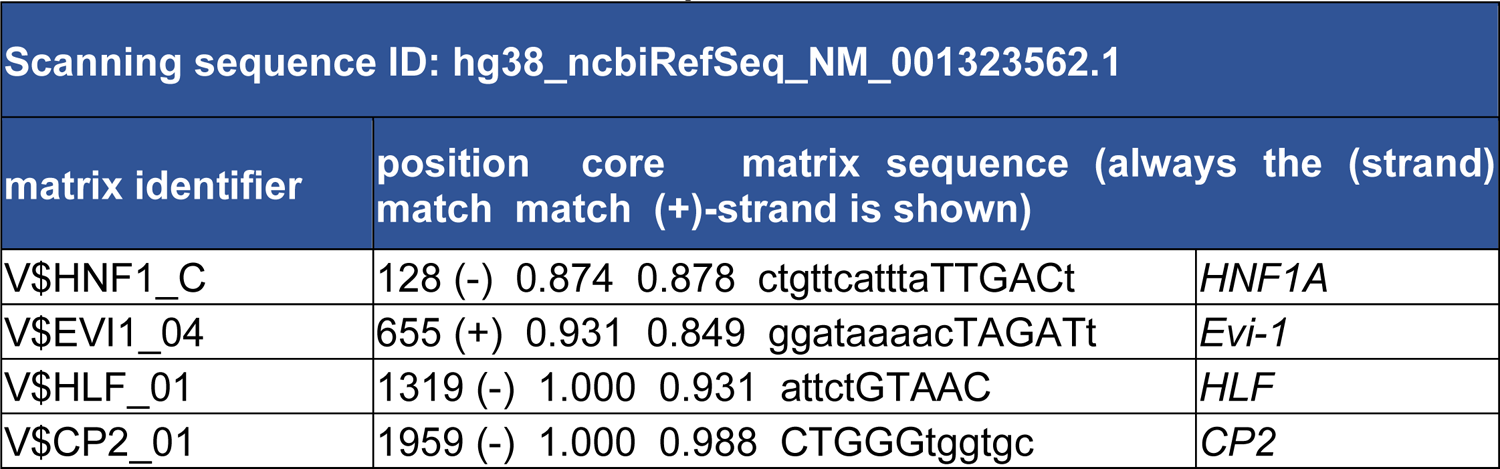
Prediction of *CRLS1* transcription factor

**table S8.**
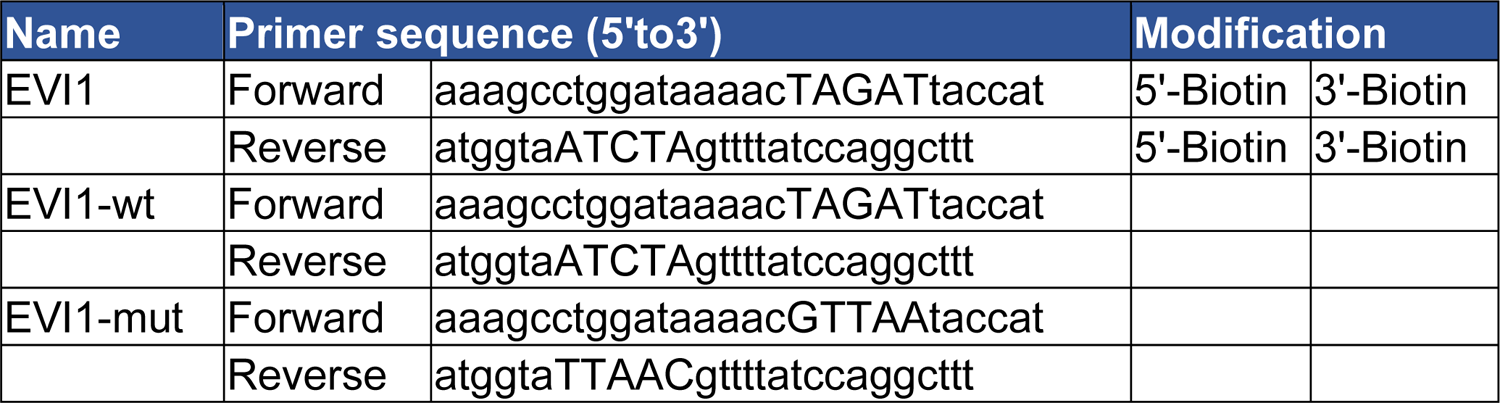
Probes used in EMSA

**table S9.**
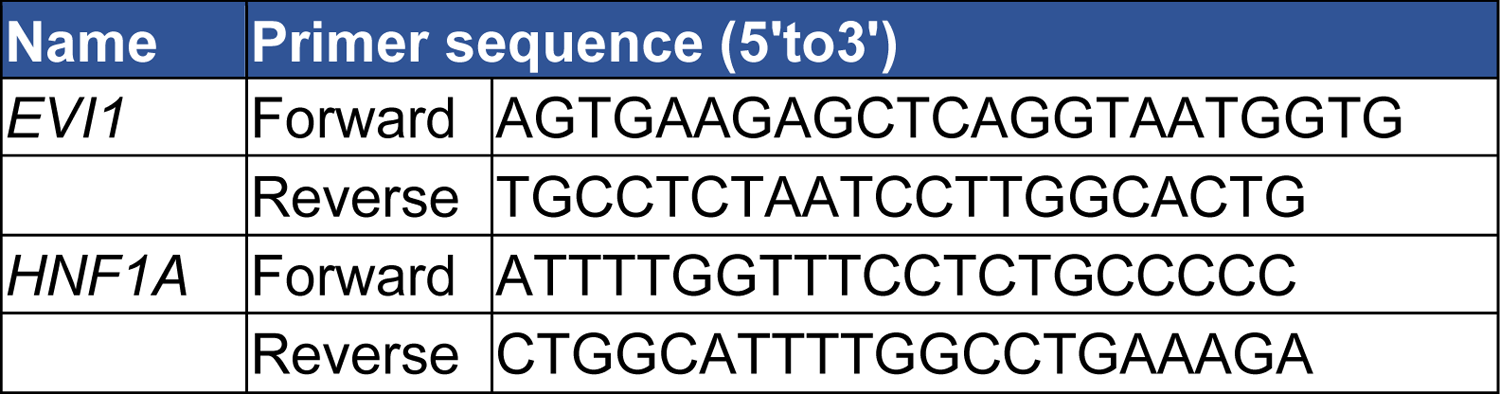
Primers used in ChIP assays

